# High mannose N-glycans on red blood cells as phagocytic ligands, mediating both sickle cell anaemia and resistance to malaria

**DOI:** 10.1101/2020.11.26.399402

**Authors:** Huan Cao, Aristotelis Antonopoulos, Sadie Henderson, Heather Wassall, John Brewin, Alanna Masson, Jenna Shepherd, Gabriela Konieczny, Bhinal Patel, Maria-Louise Williams, Adam Davie, Megan A Forrester, Lindsay Hall, Beverley Minter, Dimitris Tampakis, Michael Moss, Charlotte Lennon, Wendy Pickford, Lars Erwig, Beverley Robertson, Anne Dell, Gordon D. Brown, Heather M. Wilson, David C. Rees, Stuart M. Haslam, J. Alexandra Rowe, Robert N. Barker, Mark A. Vickers

## Abstract

In both sickle cell disease (SCD) and malaria, red blood cells (RBCs) are phagocytosed in the spleen, but receptor-ligand pairs mediating uptake have not been identified. Here, we report that patches of high mannose N-glycans (Man_5-9_GlcNAc_2_), expressed on diseased or oxidized RBC surfaces, bind the mannose receptor (CD206) on phagocytes to mediate clearance. Extravascular haemolysis in SCD correlates with high mannose glycan levels on RBCs. Infection of RBCs with *Plasmodium falciparum* expose surface mannose N-glycans on healthy RBCs, which occurred at significantly higher levels on RBCs from subjects with sickle cell trait compared to those lacking haemoglobin S. The glycans were associated with high molecular weight complexes and protease-resistant, lower molecular weight fragments containing spectrin. Recognition of surface N-linked high mannose glycans, a novel response to cellular stress, is the first molecular mechanism common to both the pathogenesis of SCD and resistance to severe malaria in sickle cell trait.

## Introduction

Sickle cell disease (SCD) comprises a group of disorders affecting over 20 million individuals and is caused by a mutation causing an amino acid substitution (E6V) in the adult haemoglobin β chain (1, 2), so that the physiological haemoglobin (Hb) A tetramer, α_2_β_2_, is replaced by the HbS tetramer α_2_β^S^_2_, which can form pathological polymers. The disease is variable, with modifiers such as high levels of the fetal β haemoglobin chain, γ, resulting in α_2_γ_2_ or α_2_ β^S^γ tetramers that terminate HbS polymers and ameliorate disease. Milder disease is also associated with compound heterozygosity of the sickle cell allele with quantitative defects in α or β chains (thalassaemias) or other haemoglobin chain variants like haemoglobin C.

SCD is characterized by a multi-system vasculopathy and haemolysis, which cause much morbidity and mortality, especially in Africa. The anaemia has been ascribed to abnormal physical properties of diseased red blood cells (RBCs), which interfere with their transit through the splenic and hepatic vasculatures, so stimulating phagocytic uptake by tissue macrophages (3). However, the observation that isolated macrophages take up SCD RBCs selectively *in vitro* (4) indicates the presence of disease-specific ligands, which remain uncharacterized. Heterozygosity for HbS, sickle cell trait (SCT), affects over 250 million individuals and is maintained in the population by conferring protection against severe malaria. The mechanism underlying this protection is not fully explained, but the mutation has long been known to prevent high levels of parasitaemia (5, 6). Yet under most conditions *in vitro*, the parasites grow equally well in SCT RBCs compared to those with normal haemoglobin (7, 8), implying that protection is due to efficient immune clearance of infected SCT RBCs, and again raising questions as to the identity of the ligands responsible for mediating phagocytosis.

## Results

### Red blood cells from patients with SCD express high mannose N-glycans on their surfaces

We postulated these putative uptake ligands might be N-linked glycans, given the prominence of the glycocalyx on RBCs and the corresponding expression of lectins as key innate receptors on macrophages (9). A survey for surface ligands using a panel of plant lectins identified two panel members that bound preferentially to SCD RBC (Fig. 1a), *Galanthus nivalis* Agglutinin (GNA) and *Narcissus pseudonarcissus* Lectin (NPL). Binding was specific (Extended Data Table 1, Extended Data Figs. 1a-b, 2a) and both lectins were noted to have similar specificities for terminal mannose residues (Extended Data Fig. 1a) (10, 11). Microscopy with fluorescent GNA lectin revealed discrete patches on the surfaces of SCD (HbSS), but not healthy (HbAA), RBCs (Fig. 1b, Extended Data Fig. 1c, d). Glycomic analysis using mass spectrometry showed that SCD RBCs express N-linked high mannose glycans, hereafter high mannose glycans (Man_5_-9GlcNAc_2_; Fig. 1c), which are known ligands for phagocytosis by macrophages (9, 12) and therefore good candidates for mediating RBC uptake. High mannose glycans are also observed in the N-glycome profiles from HbAA RBC ghosts (Fig. 1c, Extended Data Fig. 3, Extended Data File 1). The proportions of high mannose glycans with respect to whole N-glycomes were not significantly different between sickle and healthy RBC ghosts (Extended Data Fig. 1e). The marked difference between GNA binding on the cell surface of HbAA compared to HbSS RBCs is therefore not explained by the total high mannose glycan content of the ghosts.

**Figure 1:**
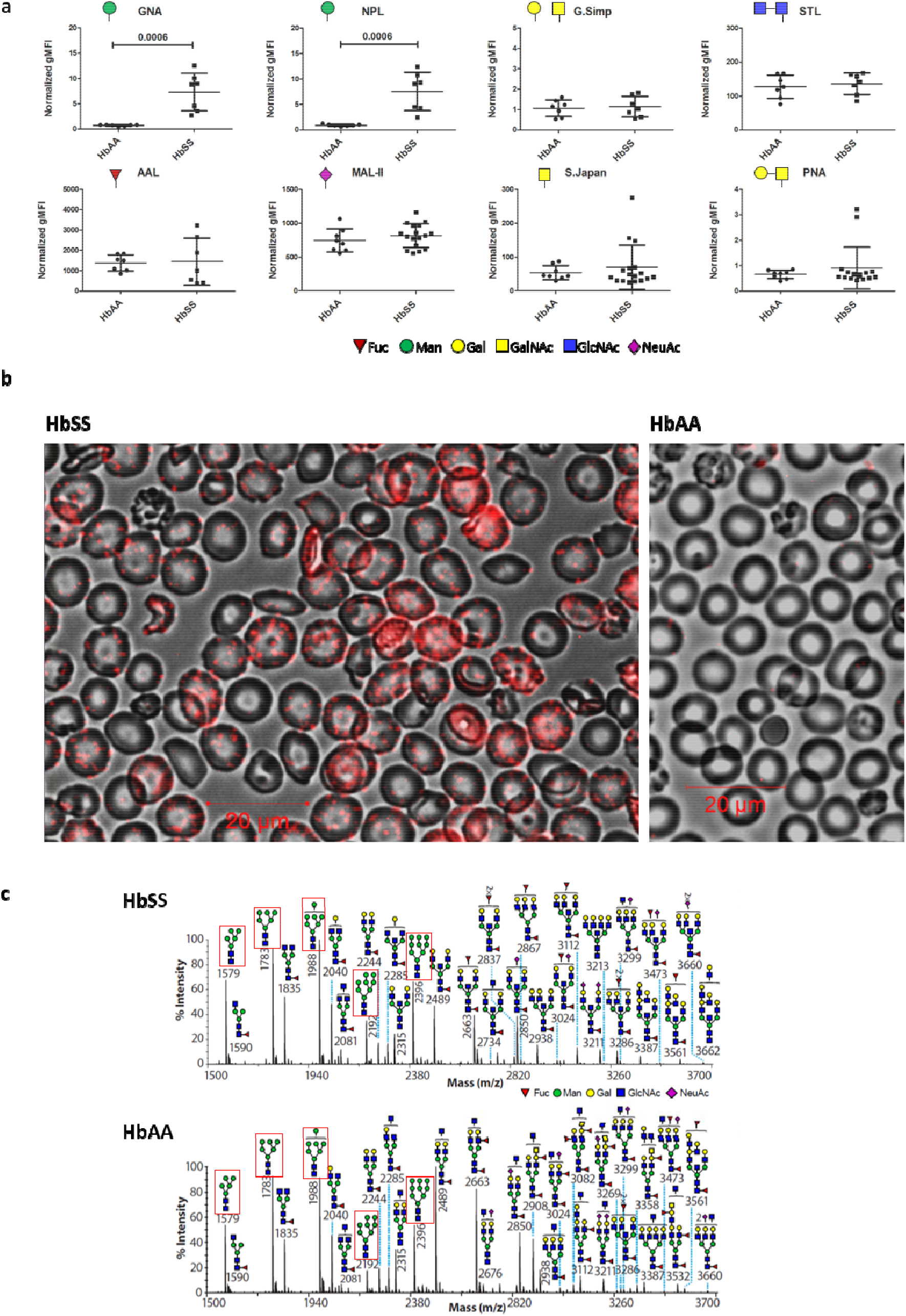
HbSS RBCs are characterized by microdomain expression of surface mannoses. a) Whole blood flow cytometry analysis of HbAA and HbSS RBCs using fluorescently labelled plant lectins, detailed in Methods. Vertical axes show normalized geometric mean fluorescence (gMFI). Symbols of terminal carbohydrate detected by plant lectins are indicated. Data shown as median +/− IQR, n=7 per group for significant differences, 2 tailed Mann-Whitney p values shown, distinct samples measured once each, 3 separate experiments. Annotation uses conventional symbols for carbohydrates in accordance with http://www.functionalglycomics.org guidelines: purple diamond, sialic acids; yellow circle, galactose; blue square, N-acetyl glucosamine; green circle, mannose; red triangle, fucose. b) GNA lectin staining (red) of HbSS and HbAA RBCs, immunofluorescence, merged with bright field. c) MALDI-ToF mass spectra (m/z versus relative intensity) for glycomic analysis of N-glycans from membrane ghosts from individual HbSS and HbAA donors. Red boxes indicate high mannose structures. Annotation uses conventional symbols for carbohydrates in accordance with http://www.functionalglycomics.org guidelines: purple diamond, sialic acids; yellow circle, galactose; blue square, N-acetyl glucosamine; green circle, mannose; red triangle, fucose. Only major structures are annotated for clarity. Full spectra of both HbSS and HbAA donors are shown in Extended Data Fig. 9.

### RBC surface mannose correlates with extravascular haemolysis in sickle cell disease

To assess the relevance of high mannose N-glycan display for RBC uptake *in vivo*, we exploited the heterogeneity of SCD arising from the interactions of HbS with other mutations in the globin loci (such as HbC, α- and β-thalassaemias) that also protect against malaria (13). If mannoses were phagocytic ligands in SCD, higher levels of mannose exposure should correlate with more severe anaemia. Despite a similar glycomic profile, RBCs from patients who were homozygous for HbS (HbSS) tended to exhibit higher binding of GNA lectin, compared to RBCs from healthy individuals containing HbAA or those with SCT (HbAS) (Fig. 2a). Patients with SCD who were compound heterozygotes for HbS and either HbC or β-thalassaemia, or who had HbSS but with mitigating α-thalassaemia or high levels of HbF, tended to exhibit low to intermediate GNA lectin binding (Fig. 2a). RBCs in other anaemias did not express high levels of exposed mannose residues (Extended Data Fig. 2a). The classical apoptotic marker for phagocytosis, phosphatidylserine, as measured by annexin V binding, was expressed at similar, low levels on RBCs from each of the clinical groups (Extended Data Fig. 2b), although it was highly expressed on positive control calcium ionophore treated, eryptotic RBCs (Extended Data Fig. 2c). Overall, GNA lectin binding correlated significantly with more severe anaemia (Fig. 2b) and other markers of haemolysis (Extended Data Fig. 2d, e), consistent with high mannose N-glycan expression driving ‘extravascular’ uptake by hepatosplenic phagocytes, which is the major mechanism of haemolysis in SCD (14). A minor, but significant, proportion of RBC loss in SCD is also accounted for by intravascular haemolysis (14). However, plasma lactate dehydrogenase (LDH) levels, a marker of intravascular haemolysis (14), did not correlate with RBC GNA lectin binding (Fig. 2c, d, e) within HbSS patients. We postulated that mannose binding lectin might bind and opsonize sickle cells, but no significant correlations between levels of this plasma protein and haemolytic phenotypes were observed (Extended Data Fig. 4a-c). Furthermore, when we added cells washed free of plasma to macrophages, SS, but not AA, RBCs were selectively taken up and this could be inhibited by mannan (Fig. 2f), indicating the macrophages expressed a receptor that interacted directly with surface mannoses.

**Figure 2:**
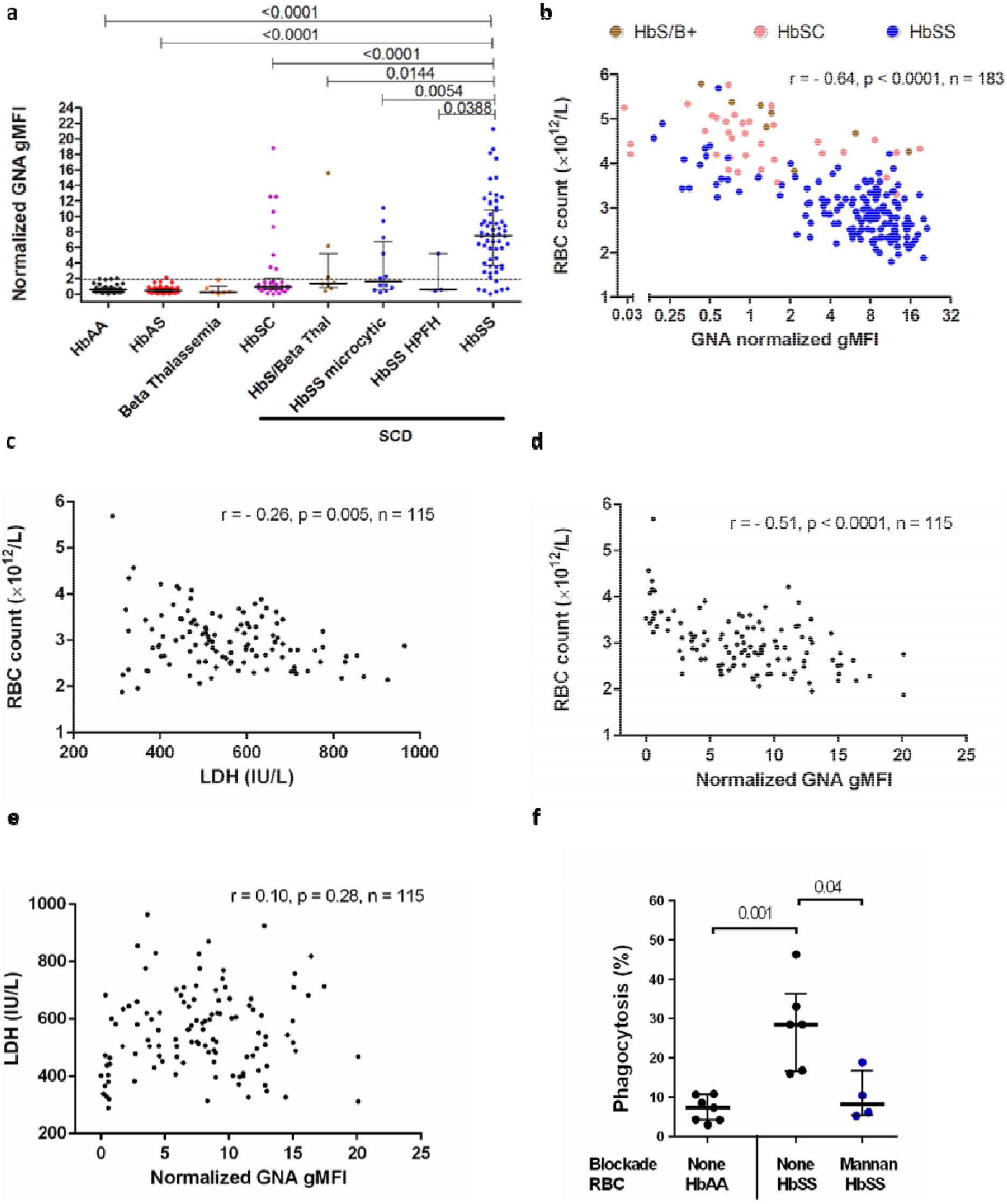
Mannose expression correlate to clinical anaemia in SCD. a) Normalized gMFI of GNA lectin staining of RBCs from peripheral blood samples, comparing RBC from patients with sub-types of SCD (milder phenotypes: HbSC (n=34), HbS/beta-thal (n=8), HbSS microcytic (n=12) and HbSS HPFH (n=3) indicate compound heterozygosity for HbC, β-thalassaemia, α-thalassaemia and hereditary persistence of fetal haemoglobin respectively) versus healthy donors (n=45), sickle cell trait (n=57) and β-thalassaemia (n=6). Dotted line indicates 90^th^ centile of GNA lectin binding within healthy samples. Data shown as median +/− IQR, 2 tailed Mann-Whitney p values shown, distinct samples measured once each, numerous experiments. b) Plot of RBC count against normalized GNA gMFI for SCD: HbS/B+ and HbSC indicates compound heterozygosity for HbS with β-thalassaemia and HbC respectively; Spearman’s rank correlation. Plots of: c) RBC count versus serum lactate dehydrogenase (LDH), d) RBC count vs GNA binding, e) LDH vs GNA lectin binding; HbSS RBCs for which corresponding serum LDH values were available; Spearman’s rank correlation, f) Percentage phagocytosis of HbAA and HbSS RBCs by human monocyte derived macrophages analysed by microscopy. Mannan inhibition as shown. Each data point represents a different RBC donor. Mann-Whitney; pooled from two experiments.

### Surface mannoses can be induced on healthy RBC by oxidative stress and are recognized by the mannose receptor (CD206)

High mannose N-glycans (Man_5-9_GlcNAc_2_) were detected in glycomic analyses of healthy (HbAA) RBCs (Fig 1c, Extended Data Fig. 3). Furthermore, permeabilization of healthy RBCs allowed GNA lectin binding in patches that colocalized with the membrane skeleton (Extended Data Fig. 5a). SCD is associated with intracellular oxidative stress (15, 16), so we determined whether exposing healthy RBCs to an oxidizing agent (Extended Data Fig. 5b) would alter the surface mannose exposure as assessed by GNA lectin binding (Fig. 3a). Under the experimental conditions applied, oxidation of healthy RBCs indeed resulted in binding of GNA lectin, with similar, but fewer, patches observed compared to unoxidized HbSS RBCs (Fig. 3b). Artefactual GNA lectin-binding resulting from permeabilization of the cell by oxidative damage was ruled out (Extended Data Fig. 5c), as was potential intracellular O-GlcNAc binding by GNA lectin (Extended Data Fig. 5d). To identify the cognate receptor on macrophages, we measured the binding of a panel of recombinant mammalian C-type lectin fusion proteins with different glycan specificities to oxidized RBCs. This survey implicated the mannose receptor (17, 18) (MR, CD206), in particular the mannose recognizing carbohydrate recognition domain (MR-CRD) (Fig. 3c). We also observed that in cultures of oxidized RBCs with human monocyte derived macrophages (HMDM) *in vitro*, uptake was restricted to MR positive cells (Fig. 3d). Furthermore, siRNA knockdown of MR in macrophages (Extended Data Fig. 6c) specifically reduced phagocytosis (Fig. 3e, Extended Data Fig. 6d); and phagocytosis was also inhibited by the competing glycans mannan or chitin, each known to block MR-CRD (19), but not inhibited by the control glycan laminarin (Fig. 3f, Extended Data Fig. 6e). Finally, MR-CRD-blocking antibody also inhibited phagocytosis of both oxidized healthy and native SCD RBC (Fig. 3g, h), which was also sensitive to mannan and chitin (Fig. 3i). Taken together, these results demonstrate that phagocytosis of mannose-displaying RBC is dependent on MR, although involvement of other receptors cannot be excluded.

**Figure 3:**
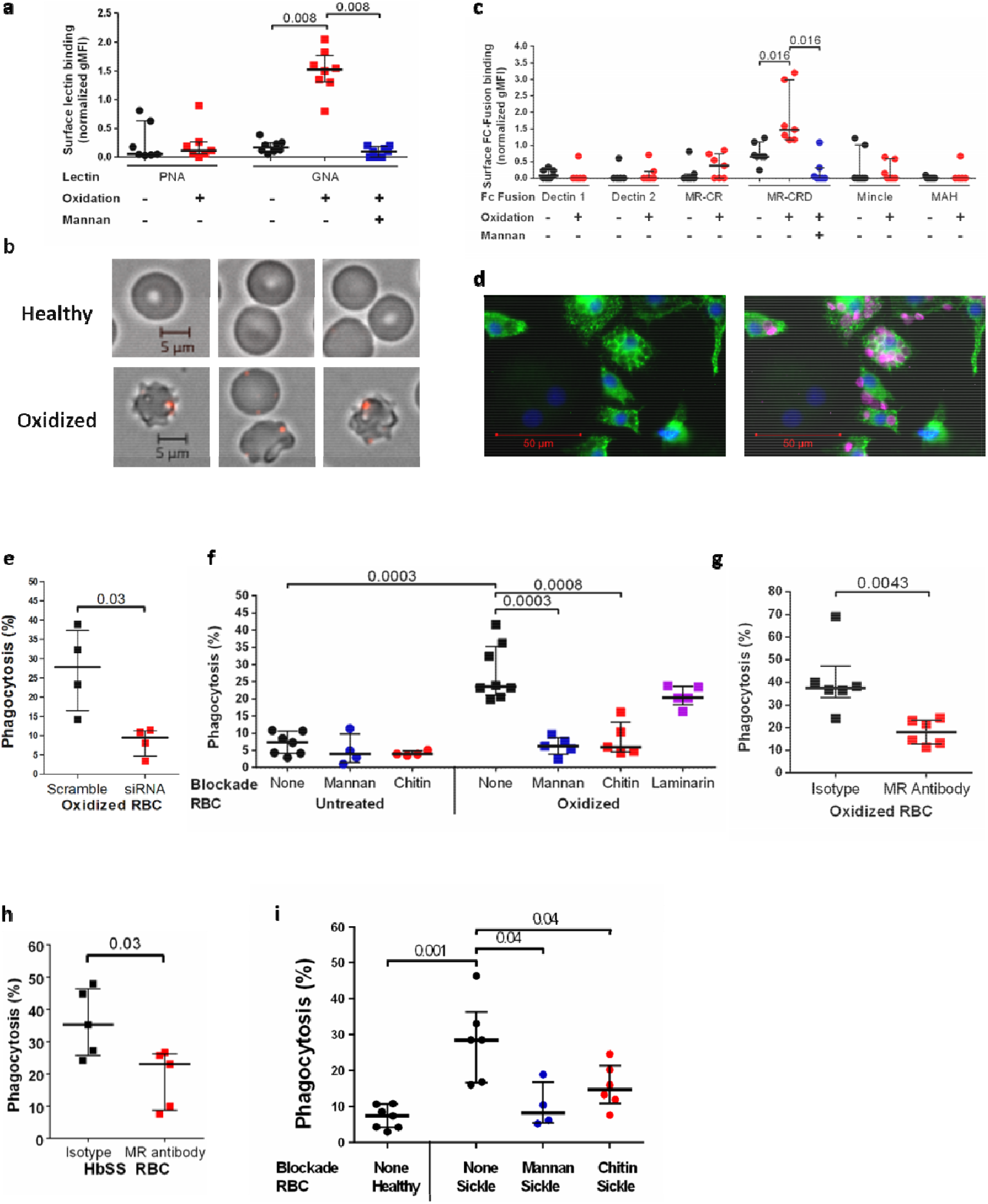
Display of membrane skeleton associated mannose patches is induced by oxidative stress and recognized by the mannose receptor on macrophages. a) PNA and GNA lectin binding to HbAA RBCs with or without oxidation. Mannan blockade for GNA lectin binding shown in blue. Wilcoxon, paired data, pooled from two independent experiments. b) Immunofluorescence microscopy of GNA lectin/streptavidin (red) staining of healthy HbAA RBCs (above) and after oxidative insult (below). c) Normalized gMFI for binding analysed by flow cytometry of murine Fc fusions with C-type lectins or sub-domains applied to oxidized versus undamaged RBCs. Mannan blockade of MR-CRD binding is shown in blue. MAH, macrophage antigen H. CR, cysteine rich. CRD, carbohydrate recognition domain. Wilcoxon, paired data, pooled from two independent experiments. d) Immunofluorescence microscopy image of human monocyte derived macrophages (HMDM) stained with DAPI (blue) and for mannose receptor (green) after incubation with oxidized HbAA RBCs, shown in magenta. e) Percentage phagocytosis of oxidized RBCs by HMDM treated with human MR specific or scrambled siRNA. Mann-Whitney, 2 experiments. f) Percentage phagocytosis of healthy or oxidized HbAA RBCs by HMDM with or without pre-blocking by mannan, chitin or laminarin g) As (f) but oxidized RBCs are blocked by MR-CRD blocking antibody 15.2 as indicated. Mann-Whitney. h) Percentage phagocytosis of HbSS RBCs with or without pre-blocking by MR-CRD blocking antibody 15.2. Mann-Whitney, 3 experiments. i) HbAA unblocked and HbSS unblocked or mannan and chitin blockade phagocytosis experiments as shown.

### GNA lectin binding proteins comigrate with spectrin containing complexes

The above data show that high mannose sugars occur in the glycomes of both HbAA and HbSS RBC, and that these sugars can be detected on the surface of HbSS RBC and oxidized HbAA RBC by GNA lectin binding. To identify the proteins carrying the high mannose sugars, extracts of HbAA and HbSS RBC membranes were analysed by western blots probed with GNA lectin. A GNA-binding doublet around 260kDa was identified in both HbAA and HbSS RBC (Fig. 4a), which is similar in molecular weight to the abundant membrane skeleton proteins α- and β- spectrin. Blots of HbSS ghosts showed additional GNA lectin-binding bands at ~160kDa, ~100kDa, ~70kDa and ~50kDa, which were not seen in fresh HbAA and HbAS RBC (Fig. 4a-b). When HbAA RBCs were stored for six weeks, to allow oxidative damage to membrane skeletal proteins (20), lower molecular weight Endo-H (N-glycan specific glycosidase) sensitive GNA lectin-binding bands corresponding in size to fragments seen in HbSS cells were seen on western blotting, and GNA lectin precipitation enriched these fragments (Extended Data Fig. 7a, b). The intensity of the 100kDa fragment was noted to correlate positively with RBC surface GNA lectin binding assessed by flow cytometry (Fig. 4b, Extended Data Fig. 7c), suggesting a role in the surface exposure of high mannose glycans. The specificity of the GNA lectin binding was confirmed by treating RBC ghosts with N-glycan specific glycosidases (PNGase F and Endo-H) prior to western blotting, which abolished GNA-binding to all of the above bands (Fig. 4c). We next attempted to determine whether treatment of RBC ghosts with N-glycanases reduced the sizes of GNA-binding bands. A molecular weight change was not observable for the high molecular weight doublet around 260kDa, although the large sizes of these proteins made minor shifts in difficult to observe. However, a ~70kDa band from HbSS ghosts did show an appropriate reduction in molecular weight, consistent with cleavage of N-glycans after treatment of RBC ghosts with PNGase F and Endo-H (Fig. 4d). Western blotting with antibodies to β-spectrin, indicated the band contained an epitope derived from spectrin.

**Figure 4:**
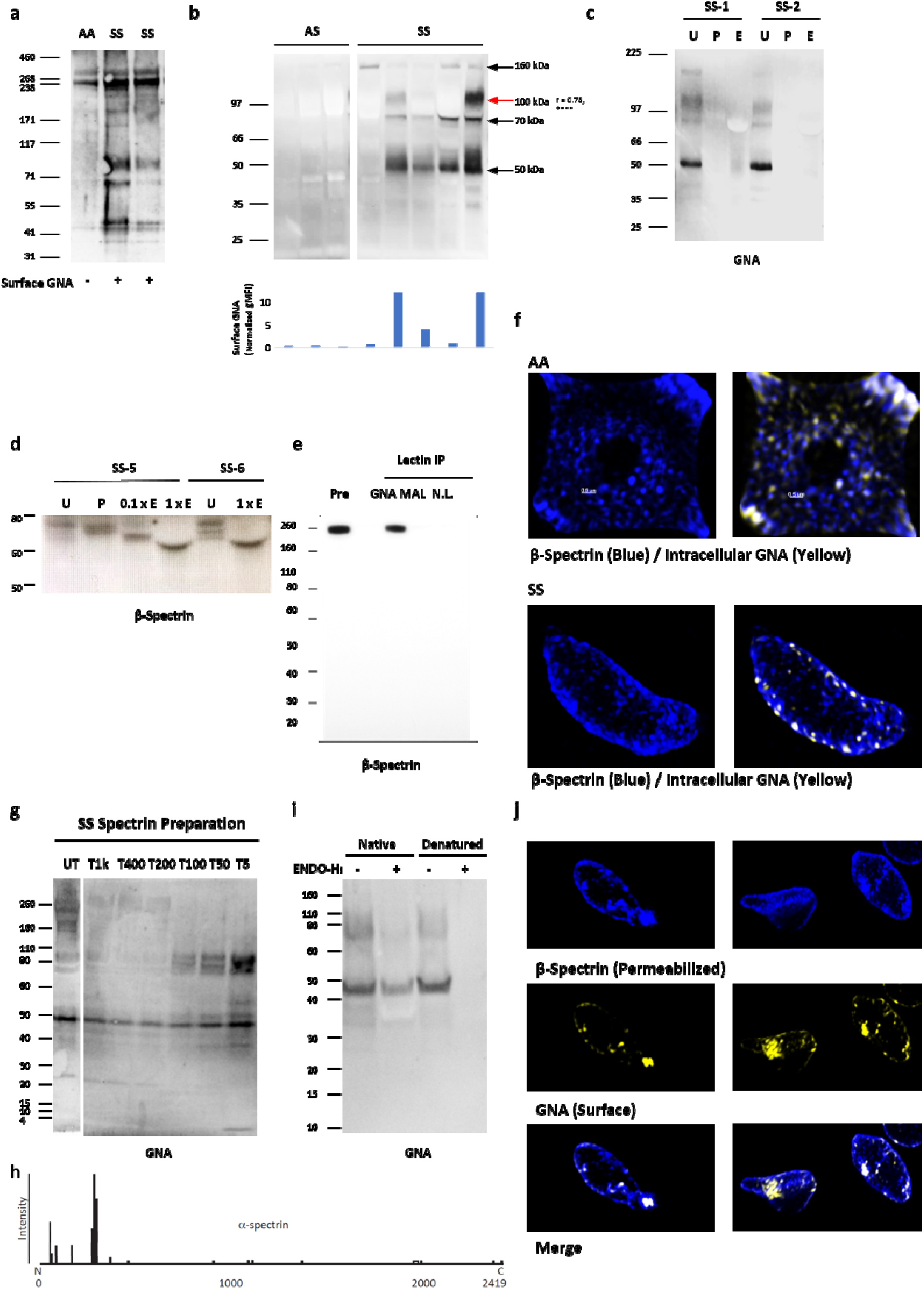
High mannose decoration of spectrin containing fragments in sickle cell disease. a) GNA lectin western blot from healthy (HbAA) and sickle (HbSS) ghosts. b) Above are shown further GNA lectin western blots from HbAA and HBSS ghosts. The histogram below the blot shows the flow cytometrically measured surface GNA lectin staining values of the RBCs used to make the ghosts, with each bar corresponding to the cells used to make the western lane above. The r value to the right of the 100kDa size label is Spearman’s rank correlation coefficient between GNA lectin staining values and band intensities, both classified ordinally as high, medium or low (n=27 measurements from 22 individuals). None of the other bands yielded significant correlation coefficients. c) GNA lectin blot from HbSS ghosts: untreated (U), treated by PNGase (P) or Endo-H (E). d) High exposure β-spectrin blot showing PNGase and partial/full (0.1X/1X) Endo-H digestion of two HbSS ghosts. e) Lectin precipitation of healthy ghosts with GNA or MAL-II lectins. No lectin control is also shown. Immunoblot with β-spectrin specific antibody. f) Super resolution microscopy image of spectrin membrane skeleton (blue) from healthy (AA) and sickle cells (SS). Yellow clusters of GNA staining are overlaid. 3D image is sliced to reveal single sheet of membrane skeleton network. g) GNA lectin blot of spectrin released from HbSS ghosts after digestion with trypsin for one hour. Untreated (UT), Tx indicates the dilution factor of trypsin relative to spectrin material. h) Peptide coverage and intensity map of α-spectrin from proteomic analysis of F40 following chymotrypsin treatment. i) GNA lectin blots showing Endo-H treatment of the 10kDa concentrate from (e) under native or denaturing conditions (urea/SDS/2-mercaptoethanol) for 24 hours. j) 3D SIM super-resolution microscopy of surface GNA lectin binding and internal β-spectrin in HbSS. HbSS RBC are first stained with GNA lectin (yellow), then permeabilized, and stained with anti-spectrin antibody (blue).

To investigate the association of spectrin with high mannose glycans species further, we carried out mass spectrometric analysis of tryptic peptides from the 260kDa GNA-binding doublet from both HbAA and HbSS RBC ghosts and found that both bands contained large quantities of α- and β-spectrin (Extended Data Table 2). As has been previously reported in RBC proteomic experiments (21), other abundant RBC proteins of lower molecular weight were also identified, including integral membrane glycoproteins such as Band 3 and Glut-1 (Extended Data Table 2). However, despite extensive mass spectrometric analyses from the 260 kDa doublet, no conventionally glycosylated peptides were identified.

Further evidence indicating covalent linkages between high mannose glycan containing glycoproteins/glycopeptides and spectrin derived peptides came from western blots of membrane extracts from HbSS RBC or HbAA ghosts treated with trypsin, which exhibited anti-spectrin binding lower molecular weight bands comigrating with GNA lectin signals (Extended Data Fig. 7c, d), particularly marked for the 50kDa GNA-binding band and α-spectrin antibodies. GNA lectin precipitation of extracts from HbAA ghosts followed by western blotting with antibodies to spectrin, detected a ~260 kDa protein (Fig. 4e). Finally, super resolution imaging of permeabilized HbAA and HbSS RBC demonstrated GNA lectin co-localising with spectrin in discrete patches scattered in the spectrin membrane skeletal network (Fig. 4f). Taken together these data support the hypothesis that spectrin containing complexes in HbSS and oxidized HbAA RBCs are N-glycosylated with high mannose glycans, although it was not possible to detect specific N-glycosylated peptides through conventional glycoproteomic approaches.

### GNA lectin binds to low molecular weight complexes that include spectrin, are protease resistant and derive from higher molecular weight aggregates

In order to generate smaller fragments of high mannose-bearing fragments that would be more amenable to characterization, HbSS ghosts were incubated with serial dilutions of trypsin, spectrin was purified from them and then probed with GNA in western blots. This showed that high concentrations of trypsin, sufficient to digest the 260kDa and 160kDa GNA lectin binding proteins, failed to degrade the ~50kDa, ~70kDa and ~100kDa GNA lectin binding gel bands (Fig. 4g and Extended Data Fig. 7e)). Indeed, the intensities of these bands, particularly that at ~100kDa, increased with higher trypsin concentrations (Fig. 4g). Prolonged, high concentration trypsin digestion eventually degraded the ~100kDa fragment, and, to some extent, the ~70kDa GNA lectin binding bands, with concurrent appearance of a new GNA lectin binding band around 40kDa, which we term F40 (Extended Data Fig. 7f). This same pattern of loss of the ~100kDa and ~70kDa GNA lectin binding fragments with concurrent appearance of F40 was also observed when HbSS erythrocytes were stored over five weeks (Extended Data Fig. 7g). Hence, protease digestion results in formation of a 40kDa protease-resistant fragment that binds GNA lectin and therefore carries high mannose glycans.

The F40 fragment was concentrated through sequential 100kDa and 10kDa cut-off concentrators (Extended Data Fig. 8a, b), and the resulting band cut out from a gel for glycoproteomic analysis. Proteomic analysis of F40 identified peptides from α-spectrin, particularly the N-terminal 370 amino acids (Fig. 4h, Extended Data Table 3). PNGase-F treatment of the purified F40 fragment released N-glycans consisting mainly of high mannoses (Man_6_-Man_9_) and complex structures (Extended Data Fig. 8c). However, once again, no specific glycopeptides could be identified. We postulated that the reason for this, and the relatively low identified protein sequence coverage, could be unconventional peptide structures arising from oxidized and glycated aggregates. Indeed, mass spectrometry confirmed the presence of lysine glycation in α-spectrin peptides from F40 at amino acids K59, K270 and K281. Furthermore, although GNA lectin binding to F40 was Endo-H sensitive, the enzyme required denaturing conditions to be effective (Fig. 4i), consistent with the possibility of protein aggregates. Additionally, the N-terminal α-spectrin antibody, B12, failed to bind to the full-length F40 band, but bound to smaller fragments after treatment with a combination of proteases, indicating a cryptic epitope (Extended Data Fig. 8d, e, f). Finally, when visualized in 3D-SIM, some HbSS cells show large aggregates of intracellular spectrin, which correspond to dense GNA lectin surface staining (Fig. 4j). Overall, our data demonstrate the existence of high mannose glycans in HbSS RBC extracts and suggest that the main GNA lectin-binding molecules are spectrin-containing glycoprotein complexes with atypical structures, including glycated forms, that make analysis by conventional glycoproteomics challenging.

### Infection of RBCs with *P. falciparum* causes exposure of high mannose N-glycans, especially those from donors with SCT

As infection of RBC with malarial parasites is associated with oxidative stress (22), we investigated whether exposure of high mannose N-glycans might be important in protection against infection with *P. falciparum*, particularly in the context of SCT. First, we determined the sensitivity of SCT RBCs to a given oxidative stress and found they bound more GNA lectin than HbAA RBC (Fig. 5a). Interestingly, the proportion of HbS correlated well with the degree of oxidative stress (Fig. 5b). These data suggested that exposure of high mannose N-glycans might contribute to the resistance of individuals with SCT to severe malaria, by enhancing clearance of infected cells. We therefore infected HbAA and HbAS RBC with *P. falciparum* and assessed high mannose N-glycan exposure as the infection progressed through ring, trophozoite and schizont stages. HbAA RBCs containing schizonts, but not trophozoites, expressed significantly higher exposed high mannose N-glycans as indicated by increased GNA lectin binding (Fig. 5c). Importantly, HbAS RBCs containing schizonts expressed even higher levels of exposed high mannose N-glycans and this increased expression extended into the trophozoite stages (Fig. 5c). *P. falciparum* infected RBCs cytoadhere to vascular endothelium, to avoid phagocytosis by hepatosplenic macrophages (23, 24), and this adhesion is mediated by the expression of PfEMP1 on late stage infected RBC (25). Reduced display of PfEMP1 on the surface of HbAS-infected RBCs is a potential mechanism of protection against malaria in SCT (26), and PfEMP1 levels show considerable variation in different HbAS donors (27). We therefore determined PfEMP1 expression in addition to mannose display and noted a marked inverse correlation (Fig. 5d).

**Figure 5:**
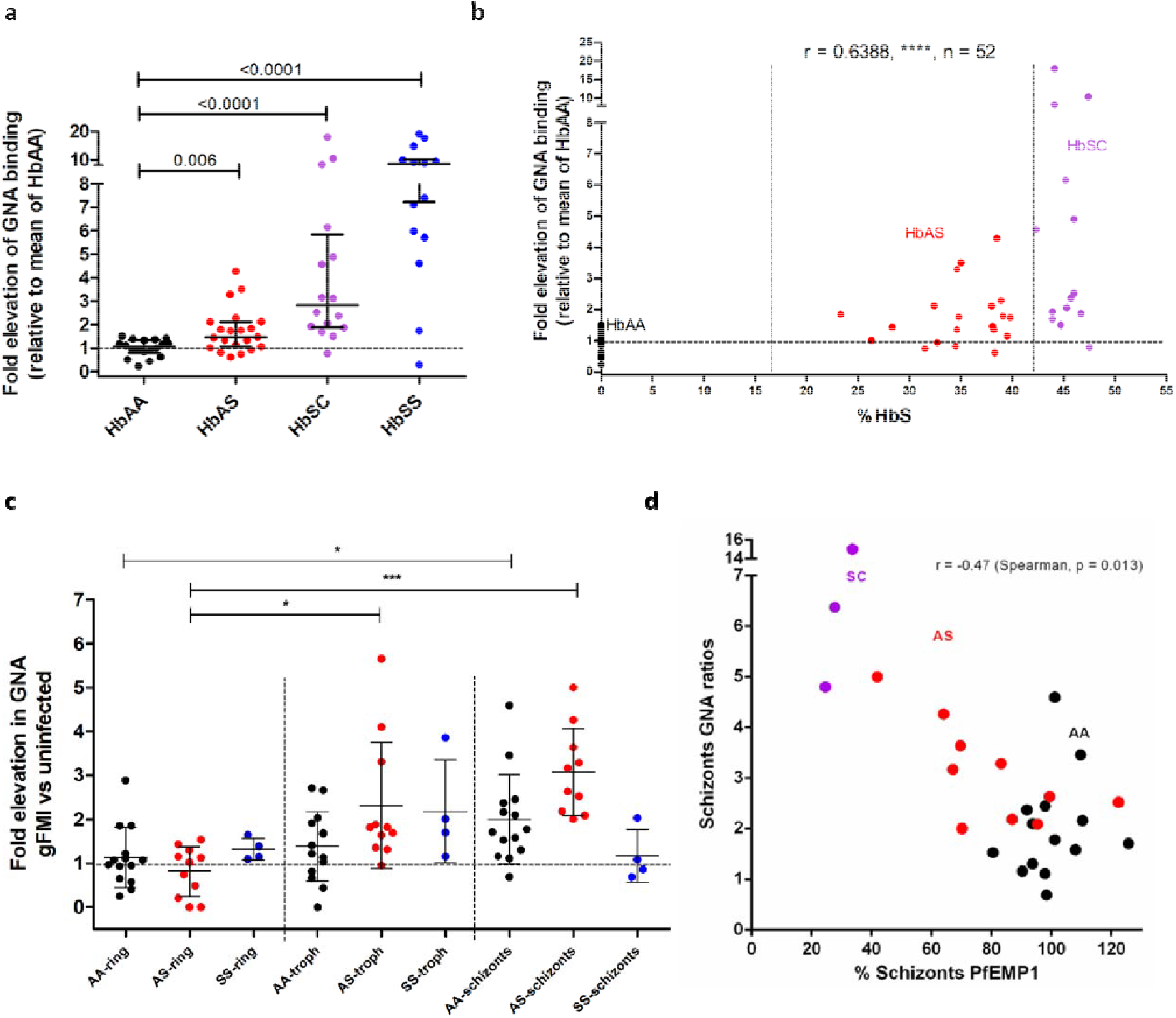
High levels of mannose are displayed by sickle cell trait RBCs in response to both oxidative stress and infection with *P. falciparum.* a) GNA lectin binding to HbAA versus HbAS, HbSC, HbSS RBCs in response to oxidative stress, expressed as ratio to mean of oxidized HbAA samples. >5 experiments, Mann-Whitney. b) HbS percentage is plotted against elevation of GNA lectin binding in response to oxidative stress as in (a). HbAA, HbAS and HbSC donor samples are shown as indicated. Spearman’s rank statistics shown. c) GNA binding to HbAA, HbAS and HbSS RBCs in response to infection with *P. falciparum*. Values for rings, trophozoites and schizonts are expressed as ratios relative to the uninfected gate. Linked ANOVA analysis for HbAA (*, p<.05), HbAS (***, p<.001), HbSS (n.s.); Tukey’s multiple comparison within each genotype, 5 experiments. d) Plot of relative GNA lectin binding for RBCs infected by schizonts against relative PfEMP1 expression. Haemoglobin phenotypes of donors as indicated, Spearman’s rank.

## Discussion

This work has identified a novel receptor-ligand pair mediating RBC clearance that underlies both the extravascular haemolysis of SCD and clearance of *P. falciparum* infected RBCs. Immunologically, the latter can be regarded as protective immunity arising from recognition of altered self, with the mannose receptor recognizing a pattern common to both diseased and infected cells. Display of high mannose N-glycans on membrane proteins could also be regarded as a new damage-associated molecular pattern (DAMP). The mannose receptor is expressed in human spleens by Lyve-1+ cells lining venous sinuses, where they form a physical barrier for blood cells to exit the red pulp and so are ideally located to perform a filtering function (28). In infection with *P. falciparum*, the parasite evades passage through the spleen by expressing adhesive proteins, notably PfEMP1, on the surface of infected RBCs, so that they adhere to endothelial cells in the systemic circulation. The inverse correlation between mannose exposure and surface PfEMP1 implies similar processes involving oxidation induced membrane skeletal rearrangements underlie both phenomena (29, 30). Reduced PfEMP1 in infected HbAS RBCs will lead to a failure of cytoadherence, and the exposed high mannose N-glycans on circulating infected RBC will induce clearance by hepatosplenic phagocytes. Together with the spleen’s role in processing high mannose N-glycan bearing RBCs in SCD, these data are consistent with spleen being the primary organ in removing RBCs infected with malaria exposing high mannose N-glycans. Our work also potentially sheds light on the reasons why those with SCD are so susceptible to infections with encapsulated bacteria, especially *Streptococcus pneumoniae*, which is the commonest cause of death in children (1). Capsular polysaccharides from pneumococcus are known to bind the carbohydrate binding domains of the mannose receptor (31). It therefore seems likely that the high mannose N-glycans on the surfaces of sickle cells would compete with bacteria for uptake by the mannose receptor.

Our work has also identified a new phenomenon whereby complexes of membrane skeletal proteins, and fragments derived from them, are associated with high mannose N-glycans, which act as an eat me signal. There are still only two accepted eat me signals in higher eukaryotes, phosphatidylserine and calreticulin (32). This work therefore adds a third ligand to perform this role. The expression of ligands for uptake attached to the membrane skeleton would allow receptors on phagocytic cells to bind to molecules with high tensile strength, which may be important for capturing cells as they transit through the spleen under conditions of high shear stress.

Protein aggregates are thought to form after attacks by free radicals, reactive oxygen and nitrogen species and glycation. These result in a wide variety of amino acid adducts (33), some of which mediate the cross-links thought to underlie the formation of aggregates. Complexes of damaged proteins, including spectrin, secondary to oxidative damage associated with haemolysis have long been recognized in RBCs (34–37), partly arising from interaction with free radicals generated by denatured haemoglobin species (hemichromes), including in SCD (38). Oxidative degradation has been shown to be one of the main causes of membrane skeletal protein alterations occurring in RBCs in storage, with proteolytic cleavage having a secondary role (20). Interestingly, spectrin containing species were detected either as low molecular weight fragments covering the N-terminus, as found in our proteomic analysis, or as high molecular aggregates (20). Oxidation during the storage period of RBCs has also been shown to inactivate glyceraldehyde 3-phosphate dehydrogenase, an important enzyme for ATP synthesis (20, 39). In turn, this leads to dissociation of spectrin from the phosphatidylserine molecules of the RBCs membrane, in an ATP dependent mechanism, resulting in increased spectrin-glycation products (40).

Therefore, either by directly acting on the cytoskeletal proteins, or indirectly through ATP dependent mechanisms, oxidative damage of RBCs is a mechanism that induces alterations in membrane protein organization leading to aggregation of membrane glycoproteins. This is in accordance with a recent report demonstrating that oxidative stress results in cluster-like structures on the membrane of RBCs as a result of possible reorganization and aggregation (41). RBCs contain various glycoproteins such as band 3, glycophorin, GLUT1, CD44 and CD47 (42). Therefore, the enhanced phagocytosis we describe could potentially be driven by the aggregation of RBC membrane glycoproteins increasing the local concentration of high mannose N-glycans, thus favouring their recognition by the mannose receptor. This is in accordance with previous reports showing that the binding affinity of the mannose receptor increases with the density of mannose-containing glycoproteins (43). Taken together, we suggest that oxidative stress in RBCs induces glycoprotein reorganization and aggregation, resulting in increased high mannose glycan bearing densities that are recognized by the mannose receptor.

In summary, we describe a mechanism whereby oxidatively-damaged, membrane protein complexes display high mannose N-glycans, which act as eat me signals important in the haemolysis of sickle cell disease and resistance against severe malaria. It therefore represents the first unified mechanism to explain both advantageous and deleterious consequences of the sickle mutation.

## Methods

### Donors

Ethical approval was obtained for the study (North of Scotland REC Number 11/NS/0026). Further samples, including donors with SCT, were obtained from the NHS Grampian Biorepository scheme (application number TR000142). Discarded anonymized samples were also obtained from patients with SCD from King’s College Hospital. HbAA, HbAS and SCD statuses were assessed by Hb HPLC (Tosoh G7). All samples from patients were collected into EDTA tubes (Becton-Dickinson).

### RBC isolation

Blood was collected into acid citrate dextrose solution tubes (ACD; 455055, Grenier) and RBC isolated by sodium metrizoate density gradient centrifugation (1.077 g/ml, Lymphoprep; 1114547 Axis-Shield). Packed RBC were diluted with an equal volume of Dulbecco’s modified Eagle’s medium (DMEM; 4.5 g/L glucose, L-glutamine; 41965, Gibco), stored in ACD (9 ml RBC/DMEM per ACD tube) at 4°C and used within 3 days unless otherwise stated.

### Lectins

Biotinylated lectins were all purchased from Vector Laboratories. They include: *Galanthus nivalis* Agglutinin (GNA, B-1245, 4μg/ml), *Narcissus pseudonarcissus* Lectin (NPL, B-1375, 4μg/ml), *Griffonia simplicifolia* Lectin II (G.Simp, B-1215, 4μg/ml), *Solanum tuberosum* Lectin (STL, B-1165, 20 ng/ml), *Aleuria aurantia* Lectin (AAL, B-1395, 33ng/ml), *Maackia amurensis* Lectin II (MAL II, B-1265, 67ng/ml), *Sophora japonica* Lectin (Vector Laboratories, no longer available, 1μg/ml). FITC conjugated Peanut agglutinin was purchased from Sigma-Aldrich (PNA, L7381-2MG, 2μg/ml).

### Flow cytometry

Whole blood flow cytometry assays were used for Fig. 1a, 2, Extended Data Fig. 1a, 2, 3b, d, e and 8a. Whole blood was washed first in phosphate buffered saline (PBS). Purified RBC were used for the other flow cytometry experiments. RBC gating was applied by forward and side scatter gating of both whole blood and purified RBC flow cytometry experiments (Extended Data Fig. 9a). Staining with anti-glycophorin A (GPA) confirmed gates contained >99% RBC (data not shown).

For lectin flow cytometry, approximately 5 × 10^6^ RBC were washed three times in PBS, and incubated for 15 minutes at room temperature in calcium buffer (10mM HEPES, 150mM NaCl_2_, 2.5mM CaCl_2_, pH 7.4) containing 10% Carbo-Free Blocking Solution (SP5040, Vector Laboratories) for whole blood flow cytometry or just buffer alone for purified RBC flow cytometry. Biotinylated lectin staining was carried out at room temperature in the same buffer as the initial blocking step. PNA-FITC and other antibody staining was carried out in PBS throughout, without contact with calcium buffer or Carbo-Free Blocking Solution. Lectin and antibody staining were carried out for 30 minutes at room temperature, protected from light. Annexin V staining was carried out according to the manufacturer’s instructions (640945, Biolegend). For biotinylated lectin staining, cells were then washed and incubated with streptavidin PE-Cy7 (0.27 μg/ml; 25431782, eBioscience) or PE (0.67 μg/ml; 554061, BD Pharmingen) for 30 minutes at room temperature. Humanized Fc fusions of murine C-type lectins (17, 44) (5 μg/ml) were incubated with RBC for 30 minutes at room temperature in calcium buffer, then detected by Alexa Fluor 647 goat anti-human secondary antibody (2 μg/ml; 109-605-098, Jackson ImmunoResearch Laboratories). In tests of their specificity for binding RBC, lectin or Fc fusion proteins were first incubated with mannan (5 mg/ml, unless otherwise stated) for 15 minutes at room temperature. Biotinylated BRIC 132 and BRIC 163 (10 μg/ml; 9458B and 9410B, International Blood Group Reference Laboratory) and anti-O-GlcNAc (1 μg/ml, RL2; 59624, Santa Cruz) binding to RBC were performed in PBS for 30 minutes at room temperature, before incubation with streptavidin secondary (for BRIC 132/163) or anti-mouse PE secondary (for anti-O-GlcNAc) for a further 30 minutes.

Prior to intracellular staining, RBC were fixed with glutaraldehyde (0.05%, 10 minutes, room temperature), permeabilized with Triton X-100 (0.1% in PBS) for 5 minutes at room temperature and then washed in PBS. Cells were washed before cytometric analysis. Data were acquired on a FACSCalibur (BD) and analysed using FlowJo v10.0 (Treestar) software. Normalized geometric mean fluorescences (gMFI) were calculated by subtracting the gMFI of secondary antibody/streptavidin-only paired controls. For PNA-FITC and Annexin V analysis, unstained controls were used for gMFI normalization.

### RBC ghost preparation

Washed RBC were subjected to ice cold hypotonic lysis in 20 mM Tris, pH 7.6, with protease inhibitor (05056489001, Roche) (45). Lysates were washed three times in hypotonic lysis buffer (37000 g, 4°C, 30 minutes) before resuspension in minimal hypotonic lysis buffer. Protein concentrations were determined by protein BCA assay (23227, Pierce). No trypsinization was performed before any glycan analysis.

### Glycomic mass spectrometry (MS)

N-linked glycan analysis from RBC ghosts were performed according to Jang-Lee *et al.* (46). MS and MS/MS data from the permethylated purified glycan fractions were acquired on a 4800 MALDI-TOF/TOF mass spectrometer (Applied Biosystems). Data were processed using Data Explorer 4.9 Software (Applied Biosystems). The processed spectra were subjected to manual assignment and annotation with the aid of a glycobioinformatics tool, GlycoWorkBench (47). Proposed assignments for the selected peaks were based on ^12^C isotopic composition together with knowledge of the biosynthetic pathways, and structures were confirmed by data obtained from MS/MS experiments.

### Proteomics

RBC ghost membranes were subjected to SDS-PAGE (10% Bis-Tris gel with MOPS running buffer). SDS-gel bands a were excised, sliced into small pieces, and destained with 200 μl of 1:1 v:v acetonitrile:ammonium bicarbonate (50 mM, pH 8.4; AMBIC). Destained gel pieces were then reduced by treatment with 10 mM DTT in AMBIC at 56 °C for 30 min, carboxymethylated in 55 mM iodoacetic acid in AMBIC in the dark at room temperature, and then subjected to overnight sequencing grade modified trypsin (Promega V5111) digestion in AMBIC at 37 °C. After enzyme inactivation (100 °C water bath, 3 min), the digested peptides were extracted twice from the gel pieces by incubating sequentially (15 min with vortexing) with 0.1% trifluoroacetic acid and 100% acetonitrile. Finally, the volume was reduced with a Speed Vac. Eluted peptides were analysed by LC–MS using a NanoAcquity UPLC™ system coupled to a Synapt™ G2-S mass spectrometer (Waters MS Technologies, Manchester, UK) in positive ion mode. 5 μL of sample was injected onto the analytical column (Waters, HSS T3, 75 μm × 150 mm, 1.8 μm). Peptides were eluted according to the following linear gradient program (A: 0.1% v/v formic acid in water, B: 0.1% v/v formic acid in acetonitrile): 0-90 min, 3-50% of B. MS data were acquired on the Synapt G2-S using a data-dependent acquisition program, calibrated using Leu-Enkephalin peptide standard. The top 20 components were selected for MS/MS acquisition. Identification of the eluted peptides was performed using ProteinLynx Global SERVER™ v3.03 (Waters) using human porcine trypsin database (Uniprot 1.0). The following were set as workflow parameters on PLGS: fixed carboxymethyl modification for cysteine, variable deamidation and oxidation modifications for glutamine and methionine respectively.

### Human monocyte-derived macrophage (HMDM) preparation and culture

Mononuclear cells were isolated by density centrifugation from whole blood and seeded at 10^6^ cells/ml in Roswell Park Memorial Institute medium (RPMI (21875-034, Gibco)), 100 U/ml penicillin, 100 μg/ml streptomycin, 292 μg/ml L-glutamine (10378-016, Gibco) and 10% heat inactivated autologous serum. Cultures were incubated at 37°C with 5% CO_2_ for 14-21 days. Cells were washed with RPMI three times prior to use.

### Phagocytosis Assay

For identification of phagocytosis by microscopy, RBC were stained with Cell Trace Far Red (CTFR) (C34564, Molecular Probes) according to the manufacturer’s instructions with minor alterations: CTFR was diluted at 1 in 500 (2 μl/ml) in RPMI with penicillin and streptomycin (RPMI/PS) media and incubated with 20 μl packed RBC for 30 minutes at 37°C, after which staining was inhibited by adding 10% FCS (10270-106, Gibco). Stained cells were washed in RPMI/PS prior to counting and addition to macrophages. RBC were added to HMDM at 5 × 10^7^ cells per well for 3 hours, before removal of cells, washing and fixation with 4% paraformaldehyde (Extended Data Fig. 6a). RBC bound, but not ingested, by HMDM were then stained with anti-glycophorin-FITC (HIR2 antibody; 306610, Biolegend). Cells were imaged using an immunofluorescence microscope (Zeiss AxioObserver Z1). Phagocytic macrophages were defined as containing at least one CTFR-positive GPA-FITC-negative RBC (determined by bright field). Three examples of oxidized RBC phagocytosis are shown in Extended Data Fig. 6b, marked ‘P’. RBC-binding macrophages were defined as associated with at least one glycophorin-FITC/CTFR double positive RBC. Analysis of HMDM phagocytosis included only the small, non-granular subset of macrophages, because of consistent association with phagocytosis and binding of RBC. For quantification of phagocytosis, 200-500 such macrophages were counted per treatment. To test specificity of HMDM recognition, the polymers mannan (10 mg/ml; M-7504, Sigma-Aldrich), chitin (50 μg/ml; C9752, Sigma-Aldrich) or laminarin (10 mg/ml; L9634, Sigma-Aldrich), or anti-CD206 blocking antibody (10 μg/ml clone 15.2 321102, BioLegend; isotype control mouse IgG1 kappa clone 107.3; 554721, BD Biosciences) were added to cultures 60 minutes before phagocytosis assays. Coumarin-stained 6 μm Fluoresbrite carboxylate microspheres, of similar size to RBC, were used to assess RBC independent phagocytosis (Extended Data Fig. 6e).

### RBC oxidation and eryptosis

Purified RBC were incubated for 60 or 30 minutes respectively with 0.2 mM copper sulphate and 5 mM ascorbic acid at 37°C in DMEM with 4.5 g/L glucose. Cells were washed in PBS 3 times prior to use. To induce eryptosis, calcium ionophore (2μM, A23187, Sigma-Aldrich) was applied at 37°C in DMEM, with 4.5 g/L glucose, to purified RBC for 3 hours.

### Reactive Oxygen Species (ROS) Production

The rate of ROS production was determined by first loading purified oxidized or untreated RBC with oxidation sensitive dye CM-H2DCFDA (10 μM; C6827, Molecular Probes) in PBS and incubating for 60 minutes in the dark at 37°C. RBC were then washed three times, resuspended in DMEM and fluorescence determined immediately by spectrofluorimeter (Fluostar Optima; BMG Labtech) with excitation of 485 nm and emission 530 nm. The rate of ROS formation was calculated for the linear portion of the fluorescence/time curve generated over six hours, which typically lasted for three hours.

### Lectin/Immuno-blotting

Ghost preparations were mixed in equal volumes with SDS sample buffer containing 8M urea (45) and heated at 100°C for 10 minutes. Ghost protein samples were fractionated by gel electrophoresis using NuPage 4-12% Bis-Tris gel (Invitrogen, NP0312BOX) and transferred by western blotting (30V, 1 hour) to polyvinylidene fluoride membrane (P 0.45 μm; 10600023 Amersham Hybond, GE Healthcare). Blots were probed with biotinylated GNA lectin (40 μg/ml; B1245, Vector Laboratories) and Streptavidin HRP (1:2500 dilution, 3999S, Cell Signalling) in calcium binding buffer (10mM HEPES, 150mM NaCl_2_, 2.5mM CaCl_2_, pH 7.4) containing 1x Carbo-Free Blocking Solution (Vector Laboratories, SP-5040) and protease inhibitor cocktail (11836145001, Roche) before development in Amersham ECL Select substrate (RPN2235, GE Healthcare). 0.1% Tween-20 was added in probing and washing steps. Loading of wells was normalized by protein concentration (~6 μg per sample). Enzymes PNGase F (P0704S, New England Biolabs) and Endo-Hf (P0703L, New England Biolabs) were used according to the manufacturer’s instructions to treat RBC ghost samples prior to electrophoresis.

### Lectin precipitation

RBC ghosts were suspended in equal volumes of calcium binding buffer containing 2% Triton X-100, pre-cleared with magnetic streptavidin beads (88816, Pierce) and incubated with biotinylated GNA lectin (1 mg/ml; B1245, Vector Laboratories), biotinylated MAL-II lectin (1 mg/ml; B1265, Vector Laboratories) or buffer only overnight at 4°C. Precipitation with magnetic streptavidin beads was performed in binding buffer and the beads washed with binding buffer containing 0.1% Triton X-100. Washed precipitates were denatured at 100°C for 10 minutes and supernatants loaded for gel electrophoresis and blotting. Blots were probed with anti-spectrin antibody (1 in 20,000 dilution; S3396, Sigma-Aldrich) and anti-mouse-HRP secondary antibody (1 in 10,000 dilution, 5887, Abcam). PBS/0.1% Tween 20 replaced calcium binding buffer for lectin blotting.

### Spectrin purification

Spectrin was purified following the method of Ungewickell *et al*. with slight modifications (48). Briefly, the ghosts were washed twice and resuspended in 3 volumes of 37°C pre-warmed sodium phosphate (0.3 mM, pH 7.2) (extraction buffer) and incubated for 20 min at 37°C. The fragmented ghosts were pelleted by centrifugation at 40000g for 1 hour at 2°C. Supernatant was used as spectrin preparation for analysis.

### Serial trypsin dilution treatment of spectrin

Spectrin preparations or ghosts were analysed for protein content by BCA assay. Titrations of trypsin at concentrations as a fraction of sample concentration were applied for one hour or longer if indicated, at 37°C. No trypsin addition was applied to the untreated sample, which was also incubated for the same duration. Samples were all heat inactivated at 100°C for 10 minutes, after diluting with 8M urea sample buffer at a 1:1 ratio.

### Chymotrypsin and pepsin digestions

Chymotrypsin and pepsin were applied at 1:5 dilution in sample. Combinatorial protease treatment over 48 hours were performed with heat inactivation for 100°C, 10 minutes at 24 hours. During pepsin treatment, sample was pre-diluted 1:1 with HCl, pH2.0. Acid was neutralized with NaOH after 24 hours, prior to addition of other proteases.

### Isolation of trypsin resistant sickle fragment (TRSF)(F40)

Approximately 20ml of HbSS ghosts, having been washed with low cold salt extraction buffer (0.3M sodium phosphate, pH 7.6), was treated with 1:5 trypsin: sample ratio overnight. Heat inactivation at 100°C was carried out for 10 minutes. Supernatant was harvested and further centrifuged to remove insoluble products. Clarified supernatant was subsequently concentrated with a 100kDa cut-off concentrator (Pierce, Thermo Fisher, 88533) and supernatant applied to and concentrated with a 10kDa cut-off concentrator to approximately 500μl.

### Mass spectrometry glycomic analysis of TRSF

Urea containing sample buffer, as above, was applied to TRSF. Coomassie bands corresponding to 40-44kD region were cut and analysed in three segments: 39-40kD, 41-42kD and 43-44kD.

### Dual colour western

Amersham western blotting machines was used to detect GNA lectin and anti-spectrin binding using Cy3 and Cy5 conjugated reagents. Data were acquired and analysed by Amersham’s inbuilt software.

### Immunofluorescence microscopy

Cell surface GNA lectin binding for immunofluorescence was performed as for flow cytometry with minor alterations (10^7^ cells per test, 8 μg/ml GNA lectin, 1 μg/ml Streptavidin PE. For Fig. 1b and Extended Data Fig.1c and 1d, 8 μg/ml GNA lectin and 1 μg/ml Streptavidin PE were pre-complexed overnight). Intracellular GNA lectin binding followed fixation (0.005% glutaraldehyde/PBS, 10 minutes, room temperature) and permeabilization (0.1% Triton-X 100/PBS, 15 minutes, room temperature). Stained cells were pulse centrifuged (≤300g) for 30 seconds (including acceleration), in 24 well, flat bottom tissue culture plates (Greiner). To stain CD206, cells were blocked for 15 minutes with 1% BSA/PBS at room temperature in the dark and incubated with Alexa-488 conjugated anti-mannose receptor antibody (1.25 μg/ml, Clone 19.2; 53-2069-47, eBiosciences). DAPI (as per manufacturer’s instructions; D1306, Thermo Fisher) staining was applied to cells post fixation/permeabilization for 30 minutes at room temperature. Cells were washed in PBS and imaged at 32x magnification using an immunofluorescence microscope (Zeiss AxioObserver Z1). Images were analysed by Zen (Black and Blue versions, Zeiss).

### siRNA knockdown of mannose receptor

Human mannose receptor (CD206) siRNA (UACUGUCGCAGGUAUCAUCCA) or a non-targeting siRNA sequence control (4390843, Life Technologies) were transfected into HMDM (RNAiMax, Life Technologies) (n = 4 donors for all siRNA experiments). Knockdown efficiency was established by determining mannose receptor expression by microscopy using CD206-Alexa-488 staining (described above) in the small non-granular macrophage sub-population by merging bright field and mannose receptor fluorescence staining. Knockdown efficiency was typically 65-85% (Extended Data Fig. 6c).

### Confocal microscopy for RBCs

For spectrin-GNA lectin double staining experiments, permeabilized RBCs were stained with anti-human spectrin antibody (1 in 50 dilution; S3396, Sigma Aldrich) concurrently with GNA lectin (8 μg/ml) in calcium buffer. Alexa Fluor 647 anti-mouse antibody (10 μg/ml; A31571, Thermo Fisher) was applied in conjunction with streptavidin PE (1 μg/ml; Beckman Dickinson) following staining of primary reagents. RBCs were gravity sedimented (30 minutes at room temperature, in the dark) onto poly-L-lysine (Sigma Aldrich) treated 8 well chamber slides (LabTek). Images were acquired by a Zeiss LSM 710 microscope.

### 3D-Structured Illumination Microscopy (3DSIM)

GNA lectin/anti-spectrin stained RBCs were gravity sedimented onto poly-L-lysine treated chamber slides (LabTek). 3DSIM images were acquired on a N-SIM (Nikon Instruments, UK) using a 100x 1.49NA lens and refractive index matched immersion oil (Nikon Instruments). Samples were imaged using a Nikon Plan Apo TIRF objective (NA 1.49, oil immersion) and an Andor DU-897X-5254 camera using 561 and 640nm laser lines. Z-step size for Z stacks was set to 0.120 μm, as required by the manufacturer’s software. For each focal plane, 15 images (5 phases, 3 angles) were captured with the NIS-Elements software. SIM image processing, reconstruction and analysis were carried out using the N-SIM module of the NIS-Element Advanced Research software. Images were checked for artefacts using SIMcheck software (http://www.micron.ox.ac.uk/software/SIMCheck.php). Images were reconstructed using NiS Elements software v4.6 (Nikon Instruments, Japan) from a Z stack comprising ≥1μm of optical sections. In all SIM image reconstructions, the Wiener and Apodization filter parameters were kept constant. Reconstructed SIM images were rendered in 3 dimensions using Imaris (Bitplane). Intracellular GNA lectin and anti-spectrin staining of healthy RBCs was performed following fixation and permeabilization. In order to co-localize surface GNA lectin with intracellular spectrin, SCD RBCs were stained with GNA lectin/streptavidin-PE staining before, and anti-spectrin/donkey anti-mouse Alexa 647 after, fixation and permeabilization.

### P. falciparum culture in RBCs of different genotypes and flow cytometry analysis

*P. falciparum* IT/FCR3 parasites were cultured at 2% haematocrit in supplemented RPMI as described (49). Mature trophozoite-infected erythrocytes were purified to >90% parasitaemia by magnetic separation with a MACS CS column (Miltenyi Biotec, Germany) (50). The purified infected erythrocytes were used to infect RBCs of different genotypes with a starting parasitaemia of approximately 0.5%. RBCs were used within 10 days post-bleed, typically 3-5 days. Cultures were gassed with 1% oxygen, 3% carbon dioxide and 96% nitrogen, then incubated at 37°C for 48-72 hours. After one cycle of invasion and growth, flow cytometry was used to assess parasitaemia and parasite maturation (51), as well as GNA lectin -binding and PfEMP1 antibody staining. This method uses internally controlled flow cytometry analysis to separate uninfected red cells, ring-stage, trophozoite and schizont stage parasites within the same culture by FACS gating, using a pair of DNA* and RNA-binding dyes (Extended Data Fig. 9b, c). Hypoxia-induced reduction in parasite invasion and growth was observed in HbAS red cells, as described by Archer *et al.* (8). However, a range of parasite stages was available within each culture to allow investigation of high mannose exposure as parasites matured through the blood stage cycle.

For flow cytometry, infected erythrocytes in binding buffer (10mM HEPES, 150mM NaCl, 2.5mM CaCl_2_, pH 7.4) were stained with Vybrant Violet (Thermo Fisher, V35003, 2.5μM) and ethidium bromide (Sigma-Aldrich, 46067-50ml-F, 1% in dH_2_0). Biotinylated GNA lectin staining was performed as described above. Streptavidin APC was used to detect biotinylated GNA lectin. A BD LSRFortessa (BD Biosciences) was used for flow cytometry and compensation between channels was carried out prior to the experiment. Relative GNA lectin binding was calculated for ring, trophozoite and schizont gates by dividing the gMFI for each gate by the value measured in the uninfected RBC gate. PfEMP1 expression was assessed using a rabbit polyclonal antibody raised against the N-terminal region (DBLαCIDR didomain) of the predominant PfEMP1 variant expressed in the culture (ITvar70, also known as AFBR6) (52), as described previously (53). The staining with PfEMP1 antibody (purified total IgG at 10 μg/ml for 30 mins, followed by APC-conjugated goat anti-rabbit Alexa 647 (Invitrogen, A21244, 2 μg/ml)) was compared to rabbit IgG control antibody (10 μg/ml, total IgG from a non-immunized rabbit, followed by secondary antibody as above). Normalized PfEMP1 gMFI was calculated by subtracting gMFI for rabbit IgG control antibody staining from that of the PfEMP1 antibody staining. Relative PfEMP1 expression for all samples is expressed as a percentage of the average HbAA schizont PfeMP1 expression per experiment, which typically contained three HbAA samples.

### Statistical analysis

Most data are treated as non-parametrically distributed and presented with medians and interquartile ranges, with the exception of Fig. 5c, where means and standard deviations are shown. Statistical significance was assessed by either two-tailed Mann-Whitney (non-paired data) or two-tailed Wilcoxon signed rank tests (paired data). Multiple comparisons between stages of RBC infection by *P. falciparum* were analysed by ANOVA. All calculations were implemented in Prism version 5.04 (GraphPad Software).

### Data Availability

The authors declare that the data supporting the findings of this study are available within the paper and its supplementary information files. Further data are available from the corresponding author upon reasonable request.

## Supplemental Information

## Acknowledgements

We are grateful for the assistance provided by both the Microscopy and Histology Core Facility, and the Iain Fraser Cytometry Centre, at the University of Aberdeen. We thank Ann Wheeler and Matt Pearson from Edinburgh Super-Resolution Imaging Consortium for technical support with 3D SIM microscopy. We also thank Janet A. Willment and Bernard Kerscher, supervised by G.D.B., for providing the Fc fusion proteins, Jeanette A. Wagener, supervised by Neil A.R.G. Gow, for providing high purity chitin, Jan Westland for obtaining blood samples and Paul Crocker for useful discussions.

Principal funding for this project was provided by Wellcome Trust grant 094847 (R.N.B, L.P.E, M.A.V). In addition, support was provided by Biotechnology and Biological Sciences Research Council grants BBF0083091 (A.D. and S.M.H.) and BBK0161641 (A.D. and S.M.H.), Wellcome Trust grant 082098 (A.D.), Wellcome Trust grants 97377, 102705 (G.D.B) and funding for the MRC Centre for Medical Mycology at the University of Aberdeen MR/N006364/1 (G.D.B).

## Author contributions

H.C. carried out experiments, analysed data and wrote the paper. S.H., A.M., B.P., J.S., H.W., M.A.F., E.B., S.L., G.K., B.M., M-L.W., A. Davie, D.T., M.M., L.H., C.L., W.P. carried out experiments. J.B., supervised by D.C.R., and B.R. obtained blood samples. A.A. obtained and analysed glycomic and proteomic data, supervised by S.M.H. and A. Dell. L.E., G.D.B. and H.M.W. helped supervise the project. J.A.R. carried out experiments, analysed data and wrote the paper. R.N.B. and M.A.V. conceived and supervised the project, and wrote the paper.

## Author information

The authors declare the existence of a financial competing interest. The University of Aberdeen has applied for patents covering diagnostic and therapeutic applications arising from the work described in this paper.

## Data Availability

Some source data are provided in the online version of the paper. The other datasets generated during and/or analysed during the current study are available from the corresponding authors on reasonable request.

---

**Extended Data Table 1:** Exclusion of possible lectin binding artefacts for SCD RBCs.

| Potential artefact | Exclusion |
| --- | --- |
| Non-specific binding of GNA lectin | • Wide lectin panel shows specificity (Fig.1a)<br>• NPL lectin binding highly correlated with GNA lectin binding across all samples (Extended Data Fig. 1a)<br>• Mannan blockade (Extended Data Fig. 1b)<br>• No binding of GNA lectin to non-SCD haemolytic anaemia, HbAA and HbAS RBCs (Fig. 2a, Extended Data Fig. 2a) |
| Non-specific binding on sickle cells | Little binding with Annexin V (Extended Data Fig. 2b), in contrast to eryptotic RBC control (Extended Data Fig. 2c) |
| Sickle cells permeable to lectins | Lack of binding of other lectins or antibodies against intracellular antigens, BRIC-132/163 (Extended Data Fig. 4d) |
| O-GlcNAc detection instead of high mannose | No surface binding by antibody RL2, specific for O-GlcNAc (Extended Data Fig. 4e for oxidized RBCs and Extended Data Fig. 4f for HbSS RBCs) |
| Reticulocytosis | Lack of expression on non-SCD RBCs with high reticulocyte counts (Extended Data Fig. 3d, e). |
| Intravascular haemolysis | LDH independence (Fig. 2c, d, e) |

**Extended Data Table 2.** Proteomic LC-MS analysis of 260kDa band cut from SDS-PAGE of ghosts from healthy (HbAA) RBCs. Proteins included are with minimum: 99.5% probability, 3 peptides identified, and 5.0% sequence coverage. Common contaminants (e.g. keratins and trypsin) have been removed. Identified proteins (UniProt accession, entry codes and description) were sorted based on the number of sequence coverage (%Cov). MW, molecular weight in Da; PLGS score, ProteinLynx Globar Server score.

| Accession | Entry | Description | MW | PLGS score | Prob. (%) | Peptides | %Cov |
| --- | --- | --- | --- | --- | --- | --- | --- |
| P02549 | SPTA1_HUMAN | Spectrin alpha chain, erythrocytic 1 OS=Homo sapiens | 279840 | 9.9 | 100.0 | 125 | 53.5 |
| P11277 | SPTB1_HUMAN | Spectrin beta chain, erythrocytic OS=Homo sapiens | 246313 | 9.9 | 100.0 | 46 | 26.6 |
| P02730 | B3AT_HUMAN | Band 3 anion transport protein OS=Homo sapiens | 101727 | 9.9 | 100.0 | 14 | 19.0 |
| P16157 | ANK1_HUMAN | Ankyrin-1 OS=Homo sapiens | 206136 | 9.9 | 100.0 | 25 | 17.0 |
| P11166 | GTR1_HUMAN | Solute carrier family 2, facilitated glucose transporter member 1 OS=Homo sapiens | 54048 | 9.9 | 100.0 | 3 | 6.3 |

**Extended Data Table 3.**
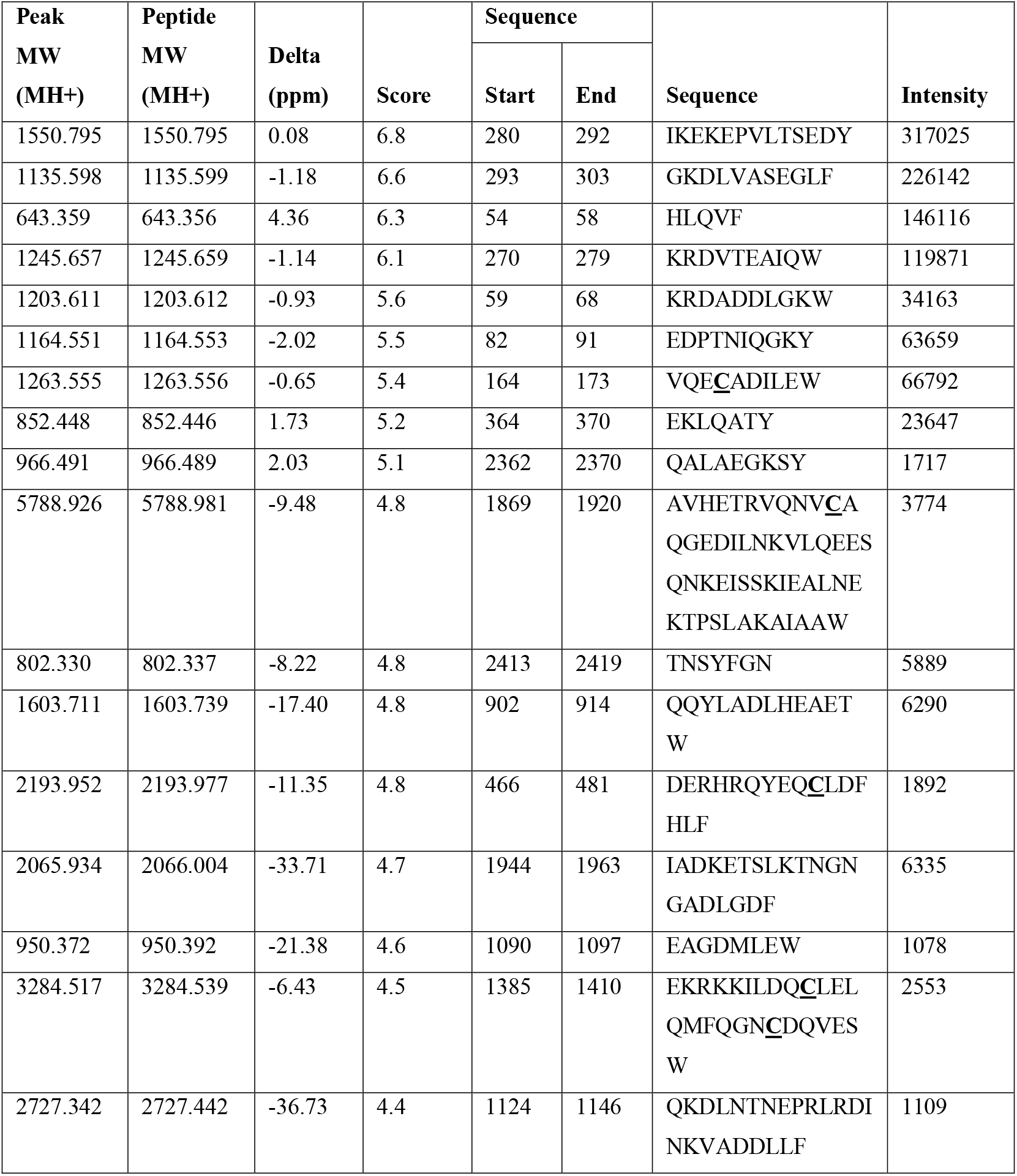
Proteomic analysis of F40 trypsin resistant gel band identified α-spectrin peptides (accession P02549, entry SPTA1_HUMAN), sorted on intensities. Peak MW, molecular weight of the protonated (MH+) peptide found in Da; peptide MW, theoretical molecular weight of the protonated (MH+) peptide in Da; delta, difference between peak MW and peptide MW in ppm; score, PLGS score. In peptide sequence, cysteine (C) amino acids in **bold** and <u>underlined</u> correspond to carboxymethylated cysteine amino acid.

| Peak MW (MH<sup>+</sup>) | Peptide MW (MH<sup>+</sup>) | Delta (ppm) | Score | Sequence |  | Sequence | Intensity |
| --- | --- | --- | --- | --- | --- | --- | --- |
|  |  |  |  | Start | End |  |  |
| 1550.795 | 1550.795 | 0.08 | 6.8 | 280 | 292 | IKEKEPVLTSEDY | 317025 |
| 1135.598 | 1135.599 | -1.18 | 6.6 | 293 | 303 | GKDLVASEGLF | 226142 |
| 643.359 | 643.356 | 4.36 | 6.3 | 54 | 58 | HLQVF | 146116 |
| 1245.657 | 1245.659 | -1.14 | 6.1 | 270 | 279 | KRDVTEAIQW | 119871 |
| 1203.611 | 1203.612 | -0.93 | 5.6 | 59 | 68 | KRDADDLGKW | 34163 |
| 1164.551 | 1164.553 | -2.02 | 5.5 | 82 | 91 | EDPTNIQGKY | 63659 |
| 1263.555 | 1263.556 | -0.65 | 5.4 | 164 | 173 | VQEC<u>AD</u>ILEW | 66792 |
| 852.448 | 852.446 | 1.73 | 5.2 | 364 | 370 | EKLQATY | 23647 |
| 966.491 | 966.489 | 2.03 | 5.1 | 2362 | 2370 | QALAEGKSY | 1717 |
| 5788.926 | 5788.981 | -9.48 | 4.8 | 1869 | 1920 | AVHETRVQNV<u>CA</u><br>QGEDILNKVLQEES<br>QNKEISSKIEALNE<br>KTPSLAKAIAAW | 3774 |
| 802.330 | 802.337 | -8.22 | 4.8 | 2413 | 2419 | TNSYFGN | 5889 |
| 1603.711 | 1603.739 | -17.40 | 4.8 | 902 | 914 | QQYLADLHEAET<br>W | 6290 |
| 2193.952 | 2193.977 | -11.35 | 4.8 | 466 | 481 | DERHRQYEQ<u>CL</u>DF<br>HLF | 1892 |
| 2065.934 | 2066.004 | -33.71 | 4.7 | 1944 | 1963 | IADKETSLKTNGN<br>GADLGDF | 6335 |
| 950.372 | 950.392 | -21.38 | 4.6 | 1090 | 1097 | EAGDMLEW | 1078 |
| 3284.517 | 3284.539 | -6.43 | 4.5 | 1385 | 1410 | EKRKKILDQ<u>C</u>LEL<br>QMFQGN<u>CD</u>QVES<br>W | 2553 |
| 2727.342 | 2727.442 | -36.73 | 4.4 | 1124 | 1146 | QKDLNTNEPRLRDI<br>NKVADDLLF | 1109 |

**Extended Data Figure 1:**
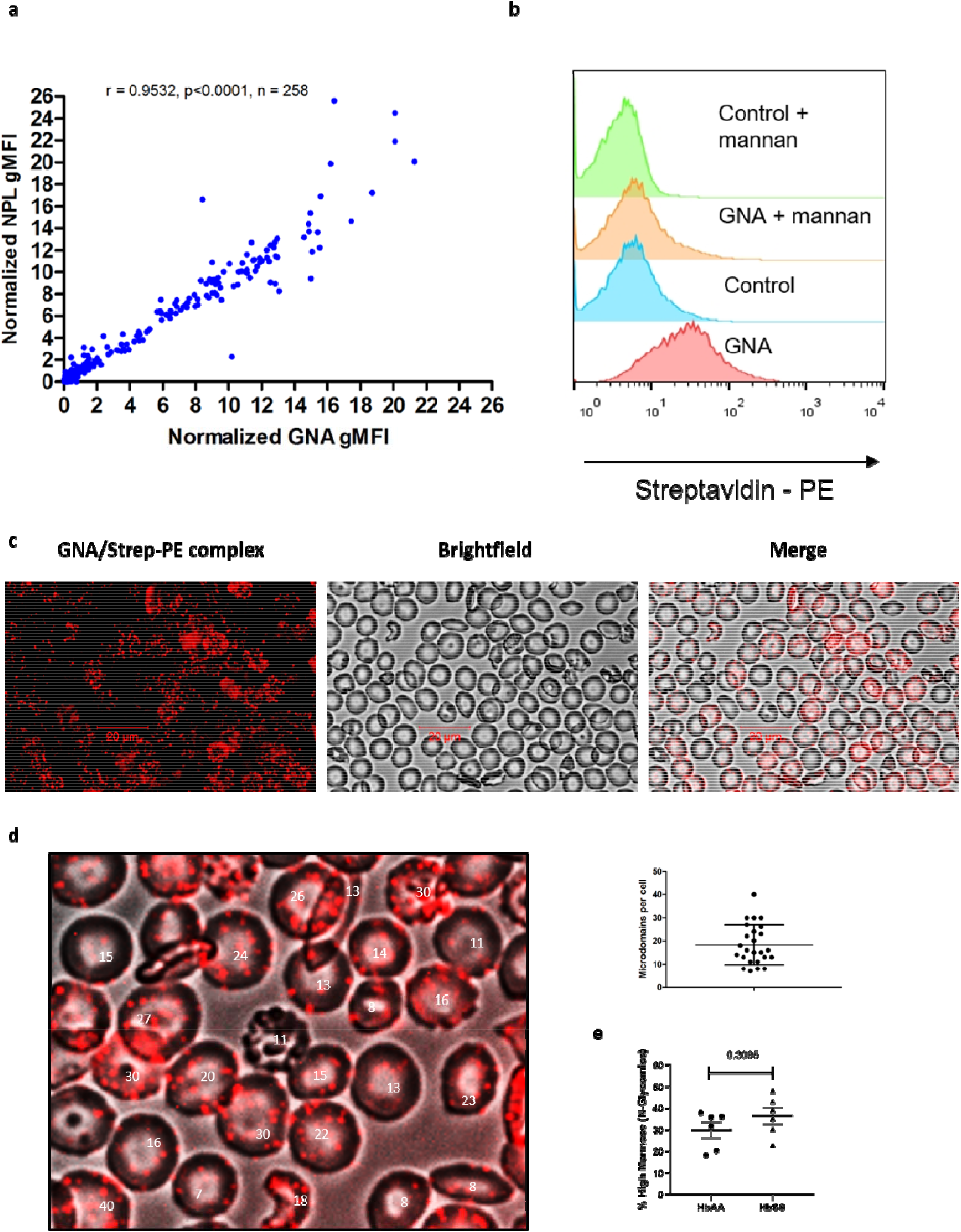
Specificity of binding of mannose binding lectins to RBCs. a) Correlation of NPL with GNA lectin surface binding; normalized gMFI, Spearman’s rank correlation. b) Flow cytometric histograms of GNA lectin and streptavidin control binding for HbSS RBCs with and without mannan blockade. Representative of 3 independent experiments from different donors. c) GNA lectin/Streptavidin PE staining of HbSS RBCs is visualized by fluorescence microscopy (PE alone, Brightfield alone and merge). d) HbSS RBCs from a section of image from c) are counted for the number of GNA lectin binding patches visualized by fluorescence microscopy (left) and plotted on the right. Mean 18.3, SD 8.6. e) Percentage high mannose structures, with respect to total N-glycans. Untreated HbAA (n = 2) and HbSS (n = 5) ghosts are analysed by N-glycome mass spectrometry. Results are pooled from four independent experiments. High mannose and complex N-glycans total 100%. Mann Whitney statistical test.

**Extended Data Figure 2:**
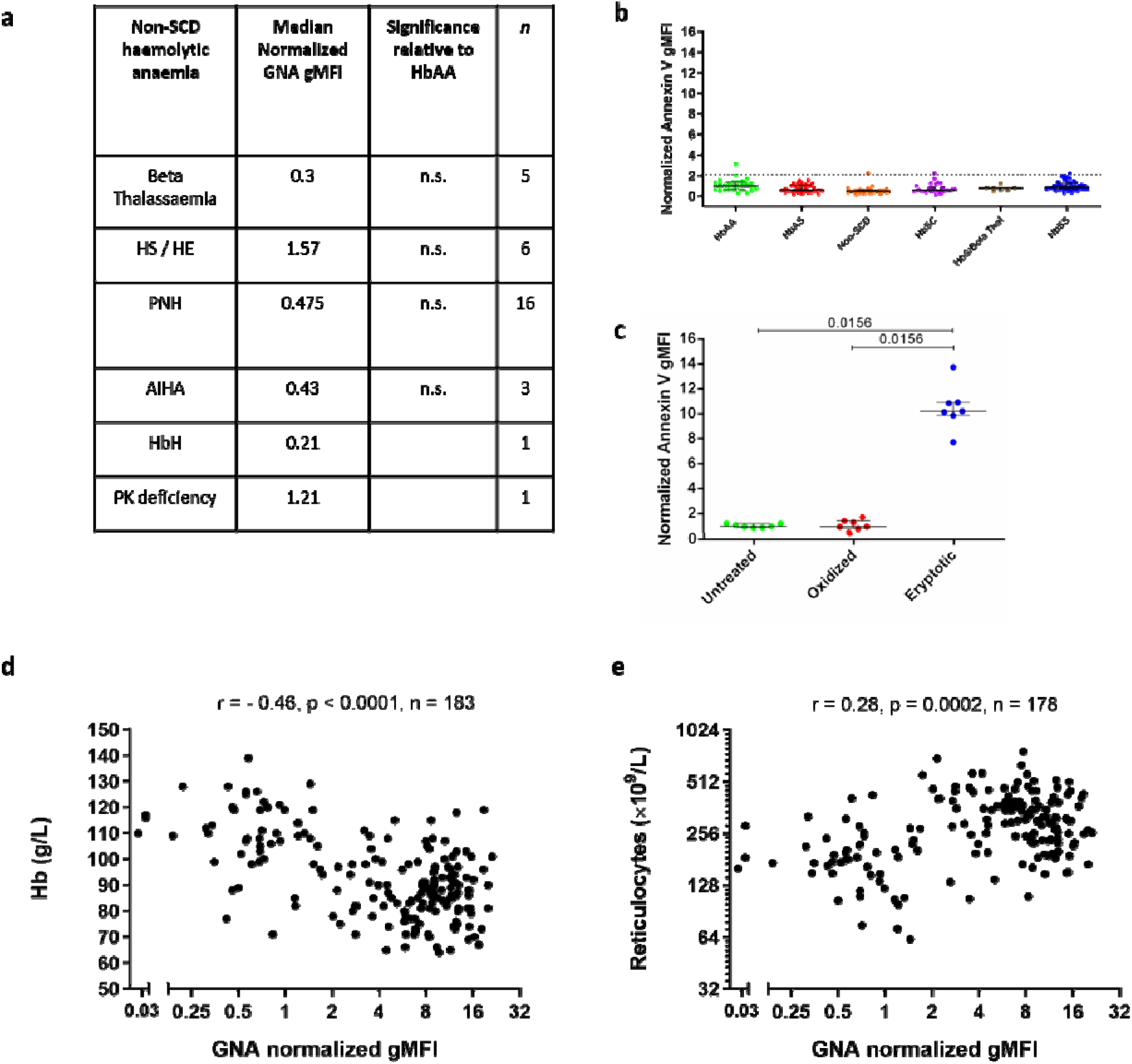
Binding of mannose binding lectins and annexin V to RBCs. a) Table of normalized gMFI values for GNA lectin binding to non-SCD haemolytic anaemias, Mann-Whitney tests relative to HbAA. b) Normalized gMFI of annexin V binding to RBCs in whole blood samples. Dotted line shows 90^th^ centile of annexin V staining within HbAA. Non-SCD include haemolytic anaemias listed in Extended Data Fig. 2a. HbAA (n=29), HbAS (n=42), Non-SCD (n=33), HbSC (n=30), HbS/Beta Thal (n=7), HbSS (n=52). No significant differences by Mann-Whitney. c) Normalized gMFI of annexin V binding to oxidized (copper sulphate/ascorbic acid), eryptotic (calcium ionophore) or untreated purified HbAA RBCs; Wilcoxon. d) Plots of haemoglobin concentrations, e) and reticulocyte counts against normalized GNA lectin gMFI for SCD including HbS/B+, HbSC (compound heterozygosity for HbS with β-thalassaemia and HbC respectively) and HbSS patients. Spearman’s rank correlation shown.

**Extended Data Figure 3.**
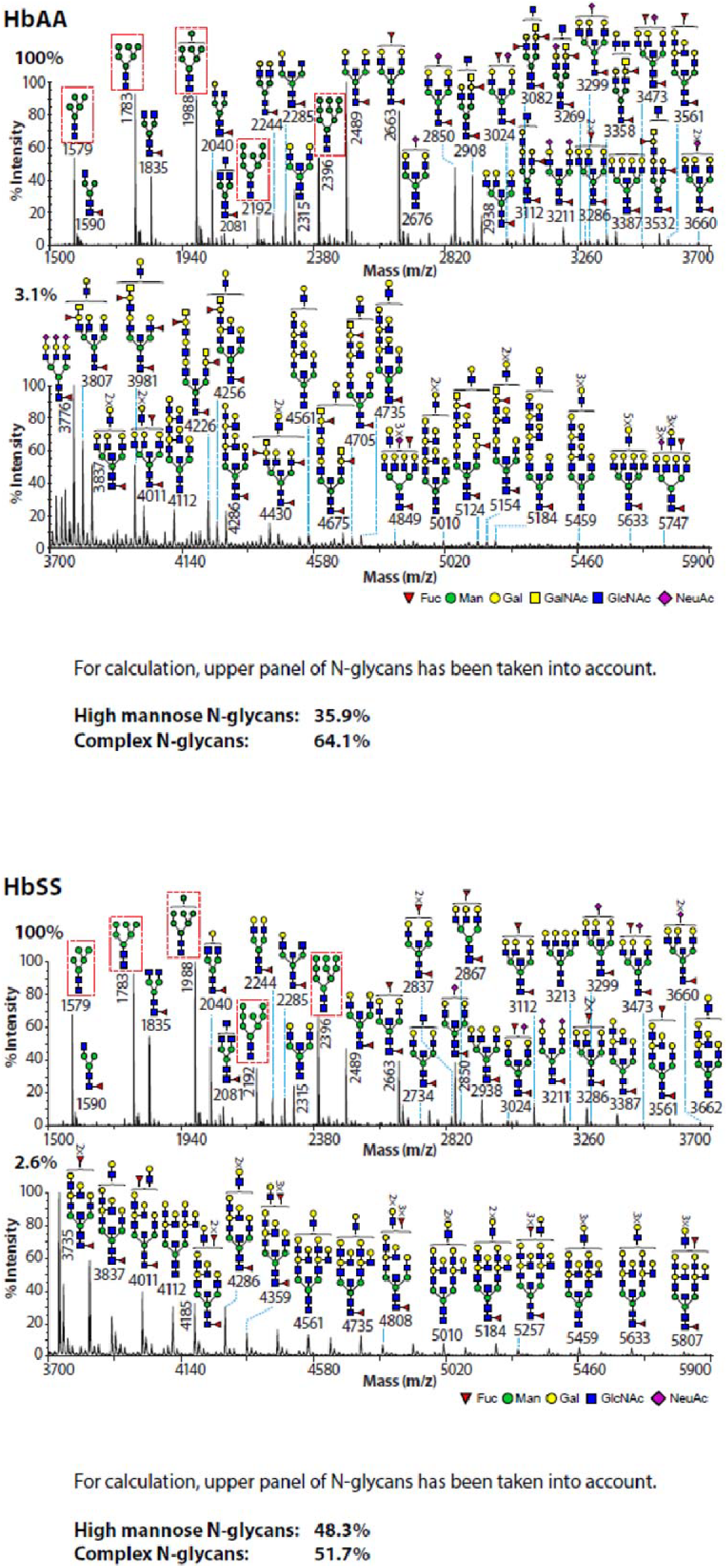
Full N-glycome spectra from HbAA and HbSS ghosts. Spectra from individual HbAA (upper panel) and HbSS (lower panel) ghosts are shown. Zoom factors are indicated by the percentages on top of the intensity axis. For calculations of percentages high mannose and complex N-glycans, the upper panel of each N-glycan profile was used.

**Extended Data Figure 4:**
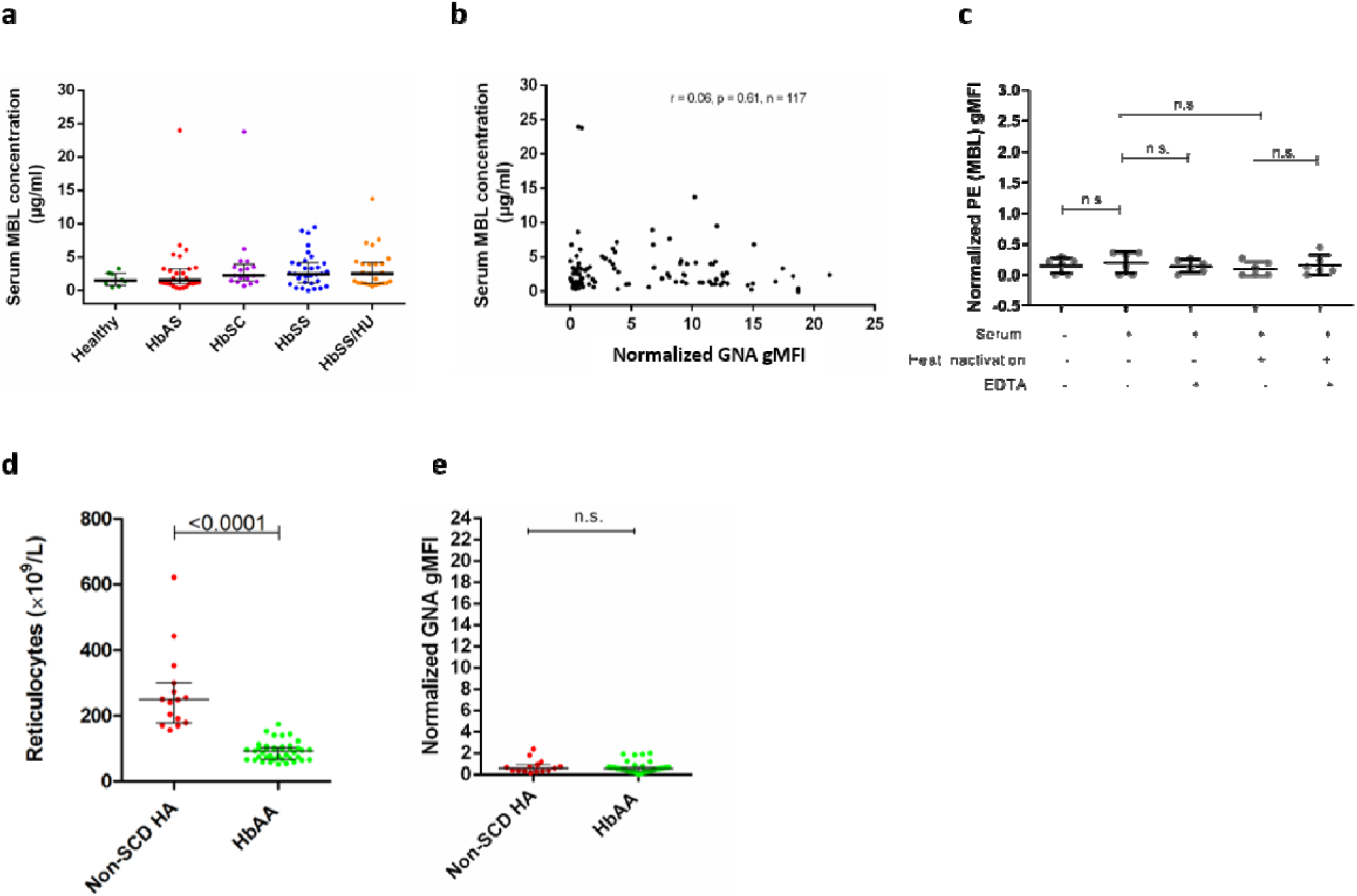
Mannose binding lectin correlations and stratification of mannose relationship to anaemia. Concentrations of serum mannose binding lectin (MBL): (a) split by clinical phenotype (HbAA (n=8); HbAS (n=27); HbSC (n=17); HbSS (not receive hydroxycarbamide treatment, n=32); HbSS/HU (receiving hydroxycarbamide treatment, n=25), (b) correlation with surface GNA lectin binding and (c) lack of direct binding to HbSS RBCs (n=6). Comparison of reticulocyte counts (d) or normalized GNA lectin binding gMFI (e) between HbAA and non-SCD haemolytic anaemias (Non-SCD HA). For d), HbAA (n=15), non-SCD (n=37). For e), HbAA (n=15), non-SCD (n=41).

**Extended Data Figure 5:**
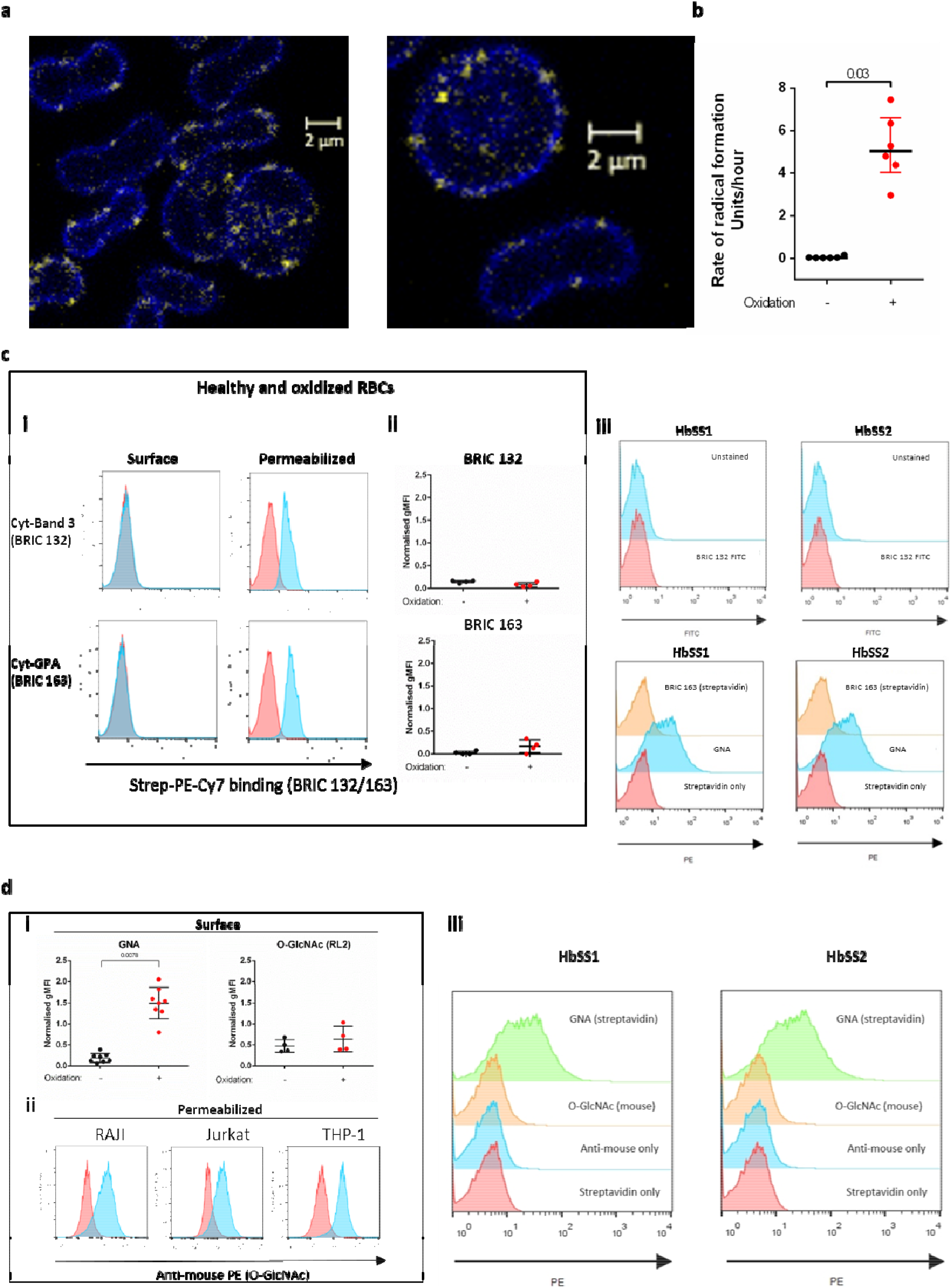
Oxidation exposes mannose on the surface of RBCs. a) Confocal microscopy of HbAA RBCs permeabilized before staining with biotinylated GNA lectin, streptavidin (yellow) and anti-spectrin (blue). b) Rate of radical formation calculated from six-hour time course measurement for healthy RBCs with or without oxidation by CuSO_4_/ascorbic acid during time course. c) Flow cytometric analysis of healthy, oxidized and HbSS RBCs using BRIC 132 (anti-cytoplasmic band 3) or BRIC 163 (anti-cytoplasmic glycophorin A). Left hand panels: i) histograms for permeabilized versus non-permeabilized binding of HbAA RBCs to BRIC antibodies. ii) BRIC antibody binding to surfaces of oxidized and undamaged HbAA RBCs. Differences not significant by Mann-Whitney. Right hand panel (iii): BRIC antibody and GNA lectin binding to RBCs from two HbSS donors, without permeabilization. d) Flow cytometric analysis of healthy, oxidized and HbSS RBC using O-GlcNAc specific antibody, RL2. i) Normalized GNA lectin and RL2 binding for undamaged and oxidized RBCs, without permeabilization. ii) Histograms for RL2 binding to permeabilized nucleated cells shown. Right hand panel (iii): RL2 antibody and GNA lectin binding to RBCs from two HbSS donors, without permeabilization.

**Extended Data Figure 6:**
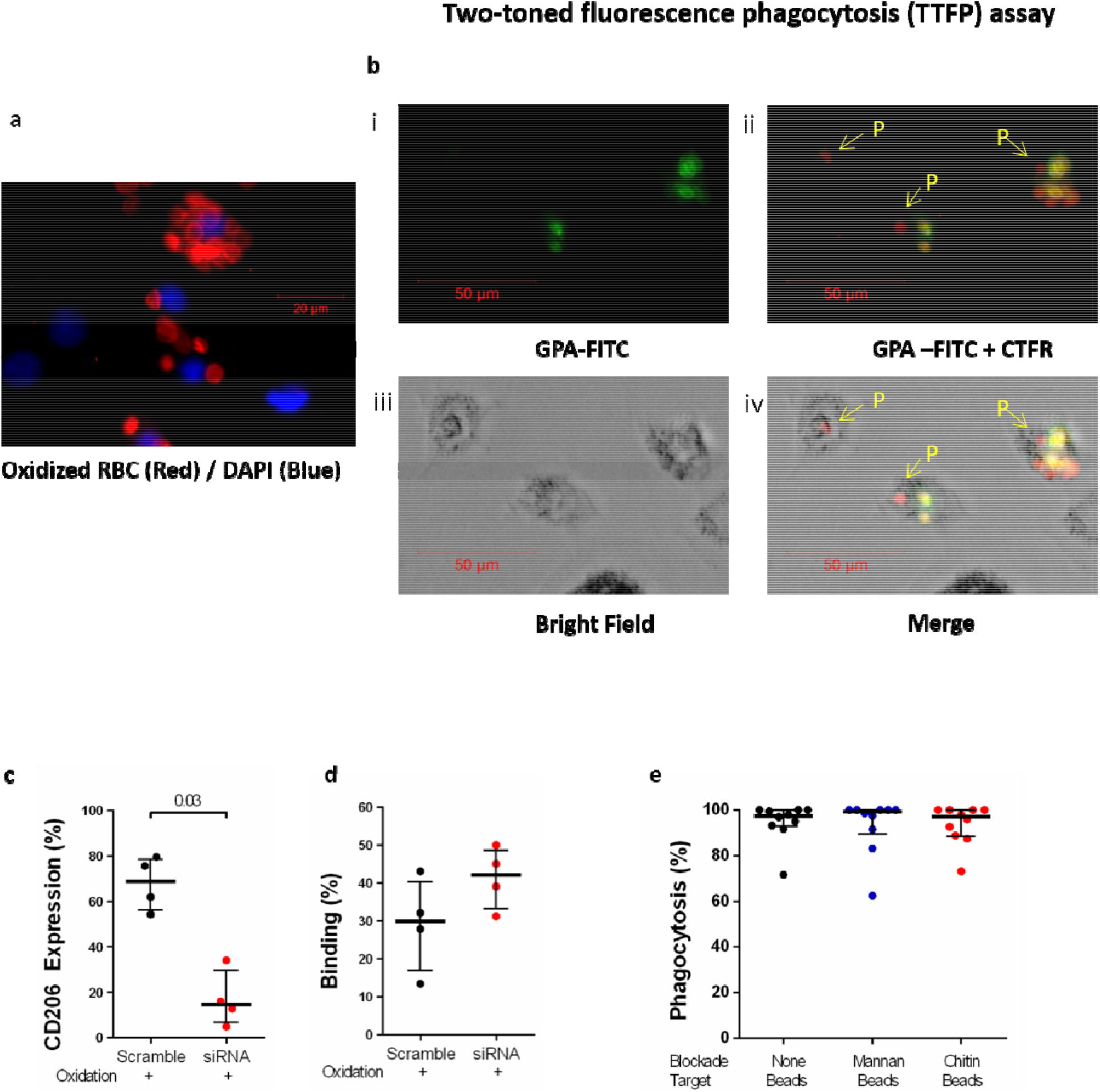
Two-toned fluorescence phagocytosis (TTFP) assay. a) Oxidized RBCs stained with Cell Trace Far Red (CTFR, red) are incubated with HMDM for 3 hours, washed with PBS, permeabilized and stained for nucleus (DAPI, blue). Immunofluorescence. Scale bar shown. b) Oxidized RBCs stained with Cell Trace Far Red (CTFR, red) are incubated with HMDM for 3 hours, washed with PBS, and counter-stained prior to immunofluorescence microscopy with anti-GPA-FITC antibody (green) to identify RBCs that are not sequestered inside macrophages: GPA-FITC only (i), GPA-FITC and CTFR (ii), bright field only (iii) and merged (iv). CTFR (red) single positive cells are counted as having been phagocytosed (letter E). Double positive (CTFR and GPA) cells are not counted as having been phagocytosed, but bound to the macrophage cell surface. Scale bar 50 μm scale as shown. c) Mannose receptor (CD206) expression assessed by microscopy for human monocyte derived macrophages treated with siRNA or scramble control. Mann-Whitney. d) Percentage surface binding of HbSS RBCs assessed by microscopy for human monocyte derived macrophages treated with siRNA or scramble control. e) Quantified phagocytosis beads by HMDM with and without glycan polymer blockade, applied in the same way and the same concentration as for RBC phagocytosis experiments.

**Extended Data Figure 7:**
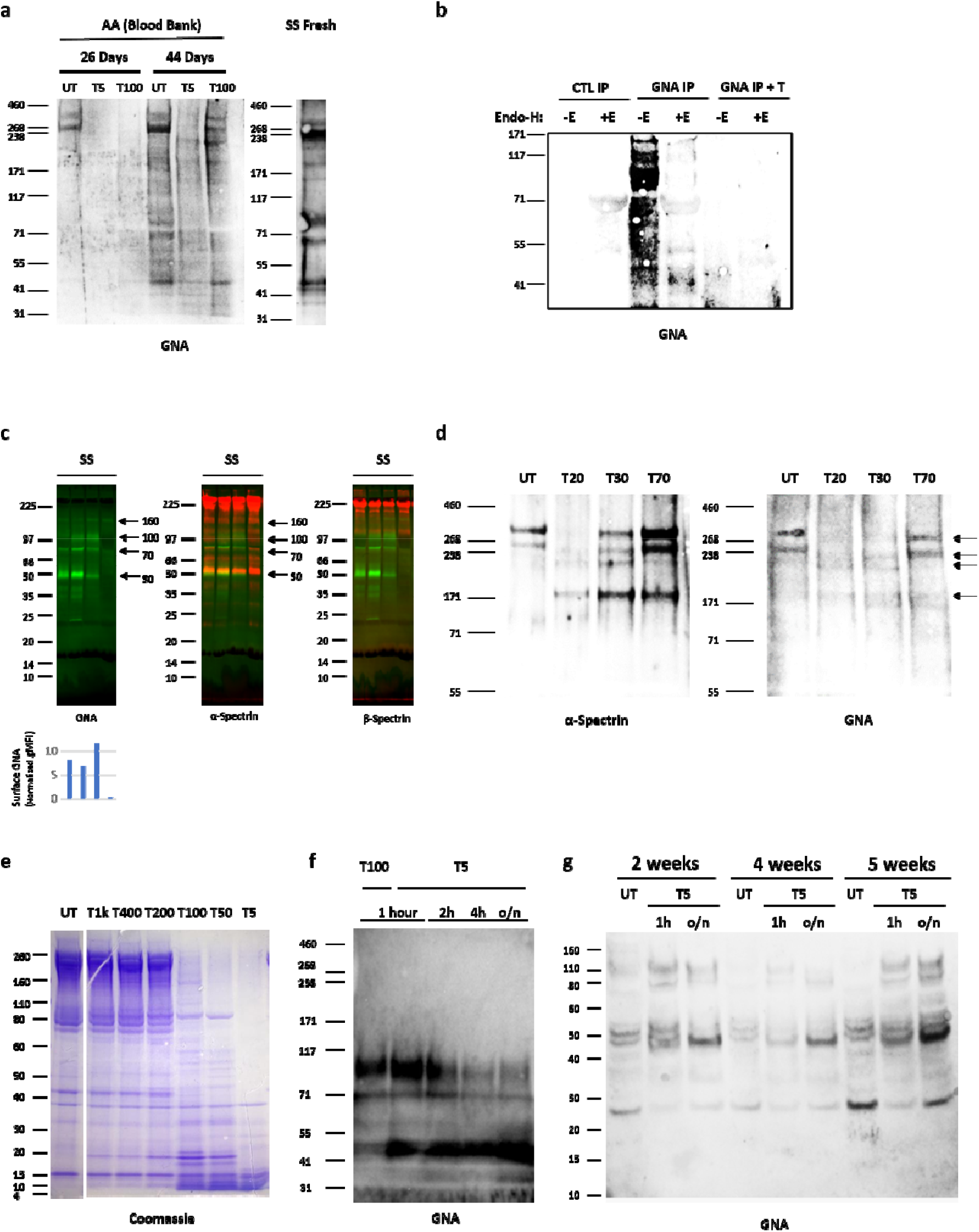
High mannose positive fragments derived from aging and proteolysis. a) GNA lectin western from HbAA ghosts isolated from erythrocytes stored from 26 to 44 days. T5 and T100 indicate trypsin digestion of ghosts, where the number indicates the dilution factor of trypsin. Comparison HbSS GNA lectin blot is shown, demonstrating correspondence in the positions of the ~160kDa, ~100kDa, ~70kDa and ~50kDa fragments. b) Stored RBC (>40 days) ghosts are Triton treated and subjected to GNA lectin precipitation or control (no lectin) precipitation. Endo-H (+E) or control digestion (−E) is applied to the eluates. Overnight trypsin treatment is also applied to a subset of GNA IP elution (GNA IP + T), prior to Endo-H or control digestions. c) Dual colour western blot from HbSS ghosts using GNA lectin (green) with either α-spectrin or β-spectrin antibodies (both red). Below are corresponding surface FACS GNA lectin staining (normalized gMFI). N.B. The 8-18% gradient gel used here does not yield bright 260kDa GNA lectin binding bands. d) Partial trypsin digestion of HbAA ghosts (T70 indicates 1:70 ratio of trypsin to sample, etc) then blotted with polyclonal α-spectrin antibody or GNA lectin. e) Coomassie stain of spectrin released from HbSS ghosts after digestion with trypsin for one hour. Untreated (UT), Tx indicates the dilution factor of trypsin relative to spectrin material. f) GNA lectin blot of HbSS ghost after prolonged trypsin treatment from 1 hour to overnight (20hours). g) GNA lectin blot of ghosts made from HbSS erythrocytes aged from 2 to 5 weeks. Untreated (UT), one hour (1h) and overnight (o/n) high concentration trypsin digestions (T5).

**Extended Data Figure 8:**
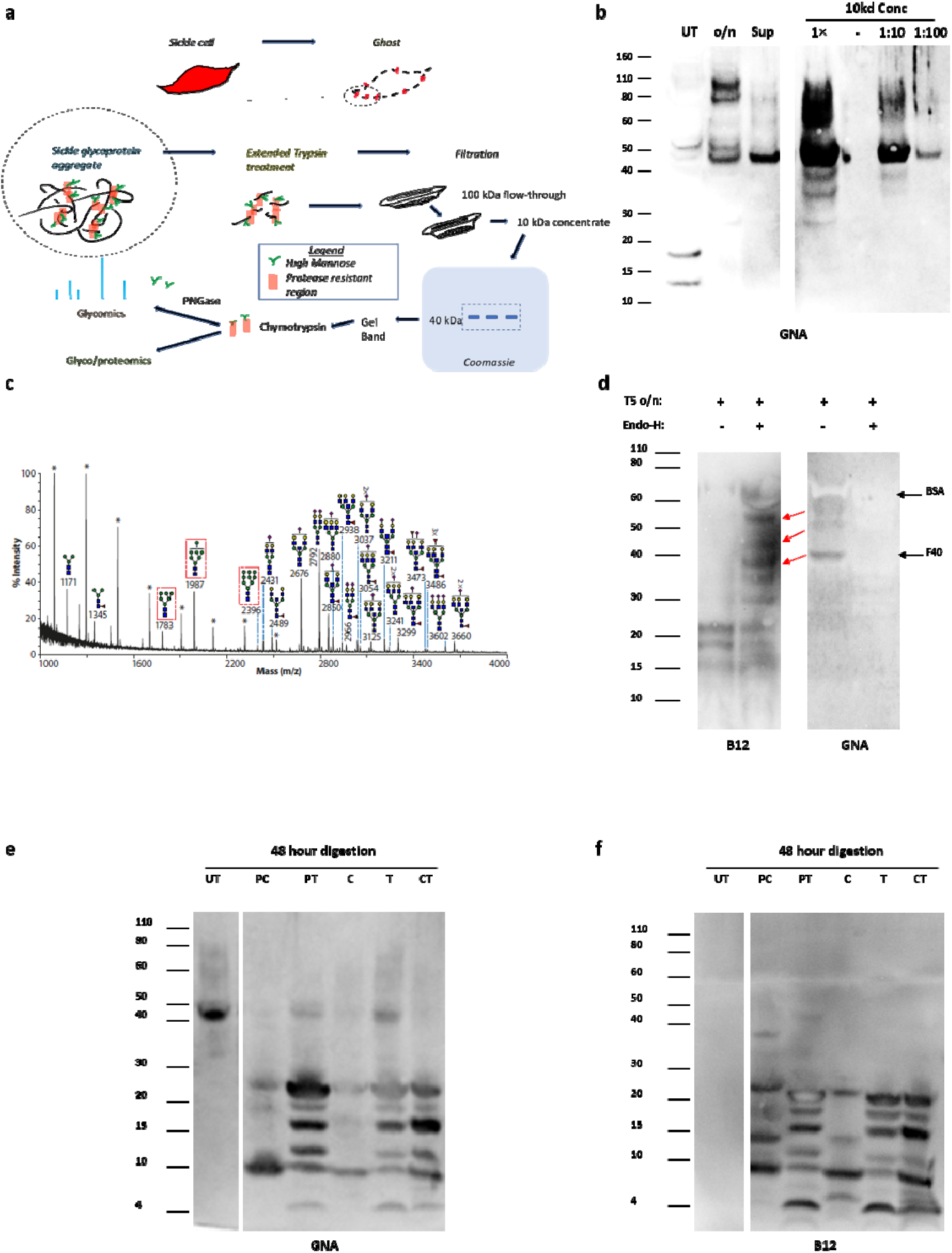
Purification and analysis of protease resistant F40 from sickle cells. a) Cartoon outlining purification process for peptide F40. Sickle cells (top right) are used to make RBC ghosts, which are then subjected to prolonged incubation with trypsin before removing the ghosts by centrifugation. The supernatant is then passed through a 100 kDa filter and concentrated on a 10 kDa filter before running on a polyacrylamide gel. The ~40 kDa band is cut out and digested with chymotrypsin (then PNGase)before analysis by mass spectroscopy for glycopeptides (and glycans). b) GNA lectin blot of enrichment process of F40 from aged HbSS erythrocyte ghosts. Untreated (UT); overnight T5 (o/n); supernatant from heat inactivated o/n sample (Sup). Final product from sequential concentration with 100kDa and 10kDa concentrators is shown as 1×, 1:10 and 1:100 dilution series. See explanatory notes below. c) Glycomic analysis following PNGase release of N-linked glycans from F40. Red boxes indicate high mannoses. Putative structures based on composition, tandem MS and biosynthetic pathways. All ions are [M+Na]^+^. Peaks annotated with an asterisk (*) do not correspond to glycan structures. Major structures are annotated for clarity. d) HbSS 10kDa concentrate (from a) was digested with trypsin (T5) overnight, heat inactivated, then digested with Endo-H or control for 24 hours. GNA lectin and B12 western blots. Red arrows show likely migration of GNA lectin and B12 reactive bands. Black arrows indicate contaminating BSA and position of F40. e) HbSS 10kDa concentrate (from a) subjected to combinatorial protease digestion in mild denaturing conditions over 48 hours. Western blot with GNA lectin. (UT) untreated, (PC) pepsin 24 hours then chymotrypsin 24 hours, (PT) pepsin 24 hours then trypsin 24 hours, (C) chymotrypsin 48 hours, (T) trypsin 48 hours, (CT) chymotrypsin 24 hours then trypsin 24 hours, with heat inactivation at 24 hours. f) As d) but with western blotting using antibody B12. **Purification and properties of F40:** F40 from overnight trypsin digestion of HbSS RBC ghosts partitions to the supernatant following heat inactivation (a). We thereby concentrated F40, through sequential 100kDa and 10kDa cut off concentrators, by approximately 50-fold (a). This preparation exhibited remarkably high GNA lectin binding, approximately 100-fold greater than full length spectrin. Sufficient F40 was purified for glycoproteomic analysis. PNGase released glycans including three high mannoses, a tri-mannose structure and other complex glycans (b). The Endo-H sensitive nature of F40 GNA lectin binding restricts the GNA lectin ligands within this pool of structures to the high mannoses: Man_6_GlcNAc_2_, Man_8_GlcNAc_2_ and Man_9_GlcNAc_2_. GNA lectin reactivity to F40 shows partial sensitivity to digestion with chymotrypsin and proteomic analysis of this digest showed the main protein represented was α-spectrin (Fig. 4i). The most abundant peptides came from the N-terminal 370 amino acids, spanning spectrin repeats 1 to 3, with only scattered, low abundance peptides identified from the rest of the molecule and none from β-spectrin. Excluding mass spectrometry incompatible regions from this region yielded 34% coverage across spectrin repeats 1 to 3, with 64% coverage within repeat 3. Identification of the N-terminal portion of α-spectrin as a major constituent of F40 from mass spectrometry analysis was also supported by western blotting using B12, an N-terminus specific α-spectrin antibody. The antibody does not bind GNA lectin reactive F40 or two higher molecular weight bands directly, but epitopes are revealed by removal of high mannoses using Endo-H (c). Furthermore, three new B12-binding bands migrated at a slightly lower positions relative to the three GNA lectin binding bands, consistent with glycosidase induced cleavages (c). 48 hour-long digestions with combinations of proteases were able to cleave F40 and, even without Endo-H treatment, unmask B12-binding epitopes (e). The GNA lectin and B12-binding fragments align remarkably well under all treatment combinations (d, e). These data suggest the high mannose decoration and unusual protein structure of F40 limits antibody access and may account for its resistance to proteases.

**Extended Data Figure 9.**
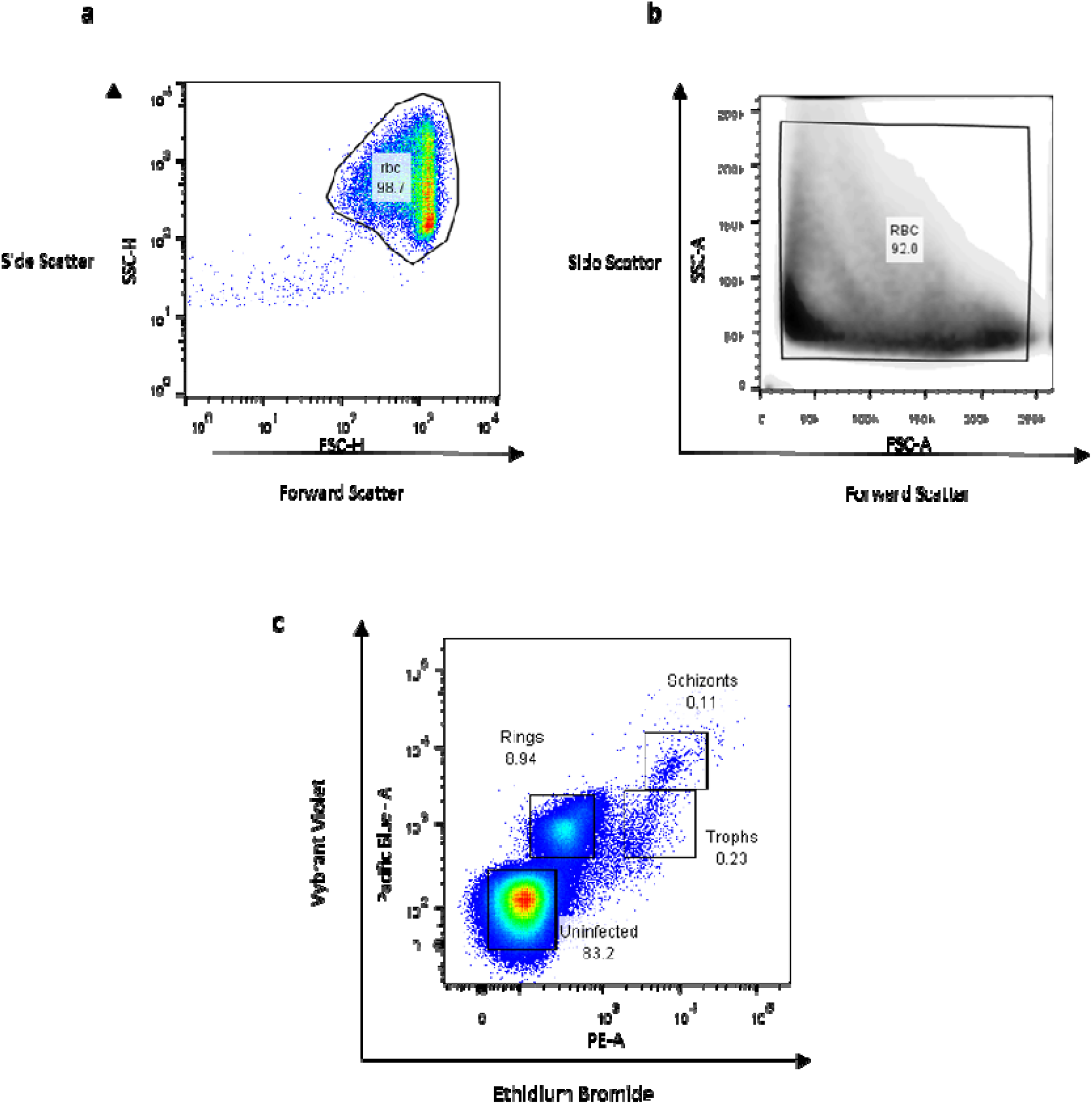
Gating strategy for flow cytometric analysis. a) Example of gating strategy for selecting whole red blood cells used in both whole blood and purified RBC flow cytometry (with the exception of *P. falciparum* infected RBC flow cytometry). b) Example of first gating of whole red blood cells used in analysis of *P. falciparum* infected RBC flow cytometry. c) Example of second gating of *P. falciparum* infected RBCs by maturation stages as defined by ethidium bromide (PE) and Vybrant Violet (Pacific Blue) staining.

